# Cyclin A2 Induces Cytokinesis in Human Adult Cardiomyocyte and Drives Reprogramming in Mice

**DOI:** 10.1101/2024.03.01.583057

**Authors:** Esmaa Bouhamida, Sangeetha Vadakke-Madathil, Prabhu Mathiyalagan, Amaresh K. Ranjan, Amir Khan, Cherrie D. Sherman, Paul E Miller, Andre Ghetti, Najah Abi-Gerges, Hina W. Chaudhry

## Abstract

Cyclin A2 (CCNA2), a master cell cycle regulator silenced in postnatal cardiomyocytes, promotes cardiac repair in animal models. However, its effect on cytokinesis in adult human cardiomyocytes remains unknown. We engineered a replication-deficient adenoviral vector encoding human CCNA2 under the cardiac Troponin T promoter and delivered it to freshly isolated cardiomyocytes from adult human hearts. Time-lapse live imaging revealed induction of complete cytokinesis with preservation of sarcomeres and calcium mobilization in redifferentiated daughter cardiomyocytes. To uncover underlying transcriptional mechanisms, single-nucleus transcriptomics of CCNA2-transgenic versus non-transgenic mouse hearts identified a cardiomyocyte subpopulation enriched for cytokinesis, proliferative, and reprogramming genes. Ultra-deep bulk RNA sequencing of adult and fetal human hearts further highlighted reprogramming pathways relevant to CCNA2-induced effects. Together, these findings demonstrate that CCNA2 can reinitiate cytokinesis in adult human cardiomyocytes and illuminate conserved molecular programs, supporting its promise as a regenerative gene therapy for the heart.

**Graphical Abstract:** 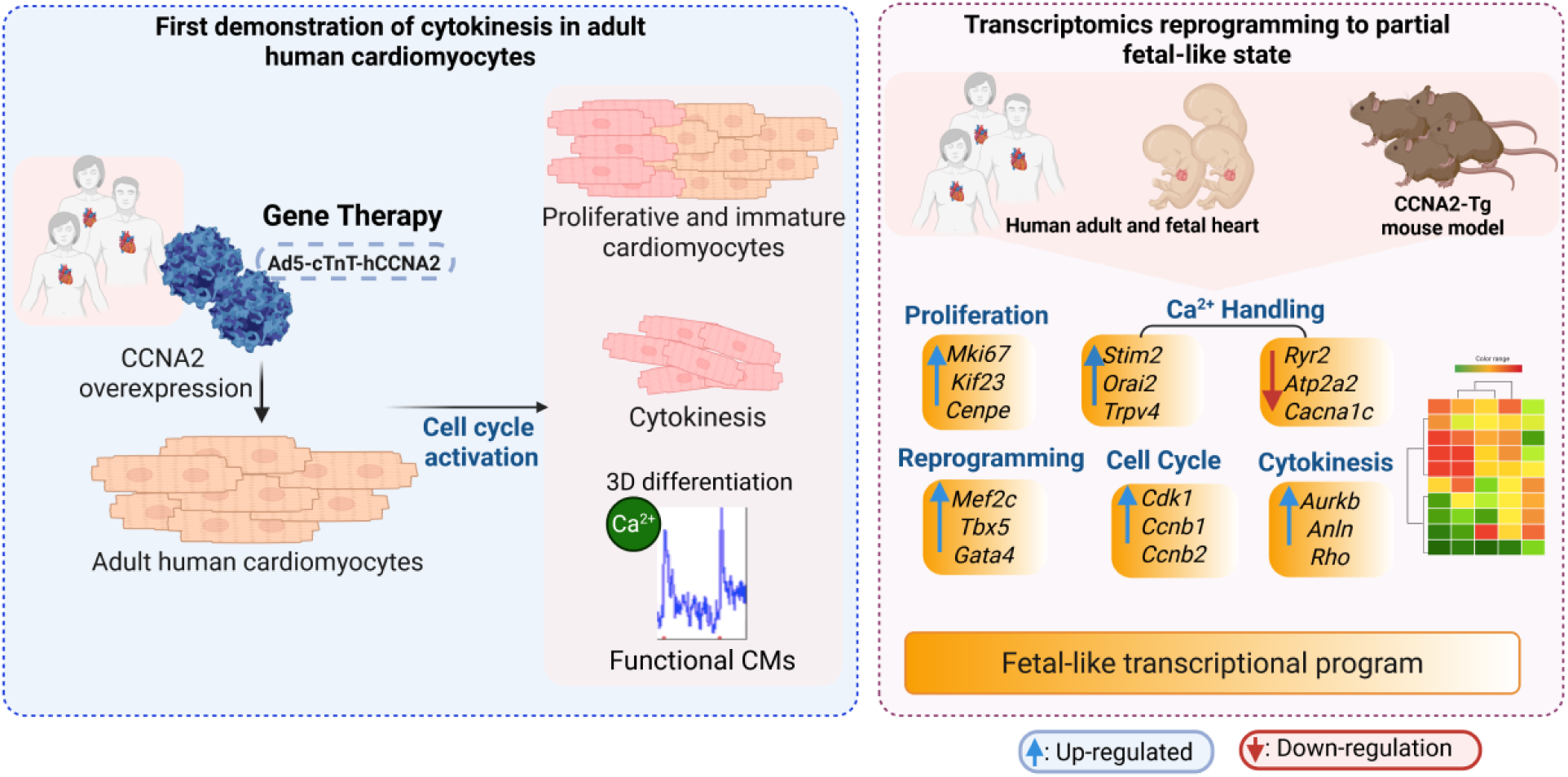

CCNA2 Induces Cytokinesis and Drives Proliferation and Reprogramming of Adult Cardiomyocytes: An Integrative Transcriptomic Analysis across Human and Mouse Models.

## Introduction

The limited proliferative capacity of adult human cardiomyocytes remains a major barrier to effective cardiac regeneration following injury, contributing to the significant morbidity and mortality associated with heart disease ^1^. Scar formation via fibrosis is, therefore, the primary response to cardiac injury. Multitudes of molecular and cellular approaches have been investigated over the past 20 years aimed at regenerating the myocardium in various states of heart disease ^2–4^. The use of stem/progenitor cell therapies for myocardial regeneration has demonstrated limited efficacy in clinical trials, without conclusive evidence of actual cardiomyocyte differentiation arising from the cell types used previously in these trials thus far. Human embryonic stem (ES) cells and induced pluripotent (iPS) cells have been evaluated in non-human primate models of heart disease with the demonstrated ability of these cell types to differentiate, at the very least, to immature cardiomyocytes with the hope that they would undergo maturation *in vivo.* However, it remains to be seen whether these approaches may be translated for human therapy ^3,5^. Beyond the differentiation and generation of new cardiomyocytes, some technical concerns still need to be addressed, such as the propensity for arrhythmias noted in primate hearts treated with ES-derived and iPS-derived cardiomyocytes ^5,6^.

There has been evidence of low-level cardiomyocyte turnover in the healthy human heart, but it is very limited, and this ability declines with age ^7,8^. Thus, cardiac regeneration in response to injury such as myocardial infarction (MI) remains a clinical challenge. Evolution may not have favored adult mammals in this regard, but certain members of metazoan species are capable of heart regeneration, and these examples may perhaps serve to enlighten us. Urodele amphibians, such as the newt, retain a remarkable capacity to replace lost anatomical tissues through epimorphic regeneration ^9^. This process relies on the local plasticity of differentiated cells near the region of injury and involves reentry to the cell cycle with loss of differentiated characteristics to generate a ‘local progenitor cell’ of restricted potentiality ^9^. The adult zebrafish heart can regenerate up to 20% of its volume via a mechanism largely dependent on the proliferation of cardiomyocytes adjacent to the area of injury ^10,11^. However, zebrafish harboring a temperature-sensitive mutation in *mps1*, a gene encoding a mitotic checkpoint kinase ^10^, cannot regenerate the heart, and scarring and fibrosis are noted in the excised areas in a manner similar to the response of the human myocardium after injury ^10,12^. In summary, the loss of a single gene can avert cardiomyocyte mitosis and mitigate the normal regenerative process, thus permitting fibrosis to proceed unimpeded.

Previously, we have reported that CCNA2 is a ‘master regulator’ of the cardiomyocyte cell cycle ^13^. Unlike other cyclins, cyclin A2 complexes with its cyclin-dependent kinase partners to regulate both key transitions of the cell cycle: G1-S and G2-M ^14–16^, and is silenced shortly after birth in mammalian cardiomyocytes ^1,17,18^. Subsequently, we have also shown that *Ccna2* mediates cardiac repair by inducing cardiomyocyte mitoses after MI in two small animal models of MI ^19–21^. As molecular mechanisms may be widely divergent across species, we performed a therapeutic efficacy study of CCNA2-mediated cardiac repair in a porcine model of MI as it closely mimics human anatomy and physiology ^22^. Therapeutic delivery of *CCNA2* one week after MI in the porcine heart induced cardiomyocyte cell cycle activation *in vivo* with marked enhancement of cardiac function as noted with multimodality imaging, including magnetic resonance imaging. The experimental pigs also exhibited significantly decreased fibrosis, lack of cardiomyocyte hypertrophy, and a 55% increase in cardiomyocyte cell numbers in the peri-infarct zones. Using the same viral vector encoding CCNA2 that was delivered *in vivo*, cultured porcine adult cardiomyocytes were induced to undergo cytokinesis with preservation of sarcomere structure and captured via live imaging ^22^.

For the purposes of clinical translation, we designed the next-generation gene therapy vector using a cardiomyocyte-specific promoter, cardiac Troponin T (cTnT), driving human CCNA2 expression in a clinical-use grade replication-deficient human adenovirus type 5. We then sought to determine whether this cTnT-CCNA2 vector would induce human adult cardiomyocytes to undergo cytokinesis. We employed a multiparametric approach combining live-cell imaging with high-resolution transcriptomics to uncover mechanistic insights underlying CCNA2-driven cytokinesis in adult cardiomyocytes. Live imaging microscopy enabled dynamic tracking of sarcomere integrity and cellular architecture during cytokinesis in adult human cardiomyocytes. To dissect the transcriptional programs and signaling pathways associated with *CCNA2* expression, we leveraged four distinct RNA sequencing datasets: (i) bulk RNA-sequencing of CCNA2-Tg and nTg cardiomyocytes, (ii) ultra-deep bulk RNA-sequencing of adult and fetal human hearts, and (iii) single-nucleus RNA sequencing (snRNA-seq, 10x Genomics). We also compared the gene expression profiles in these three datasets to previously published data regarding hypertrophic signaling pathways activated by total aortic banding (iv).

This integrative approach defined the molecular framework by which CCNA2 reactivates cell cycle networks, promotes cytokinesis, and induces a regenerative transcriptional state in postnatal cardiomyocytes, offering insights for future cardiac regenerative therapies.

## Results

### cTnT-CCNA2 adenovirus vector designed for therapeutic use induces expression of CCNA2 in cultured adult human cardiomyocytes

First, we designed a therapeutic grade human CCNA2 adenovirus vector to selectively express human cyclin A2 (CCNA2) in cardiomyocytes by cloning human cDNA (NCBI Reference Sequence: NM_001237.4; 374-1672 bp) downstream to the cTnT promoter **(Figure 1A)**. The cultured adult human cardiomyocytes were transduced with cTnT-hCCNA2 (test) versus cTnT-eGFP (control) adenoviruses with a multiplicity of infection (MOI) of 100 each for assessing the induced expression of CCNA2. We observed significantly increased expression of CCNA2 in the cultured adult human cardiomyocytes transduced with the test compared to the control adenovirus **(Figure 1B)**. We utilized cTnT-H2B-GFP to confirm that the nuclei of the daughter cells belonged to cardiomyocytes. A schematic overview of the study design and integrated analyses is provided in **Figure S1**.

**Figure 1.**
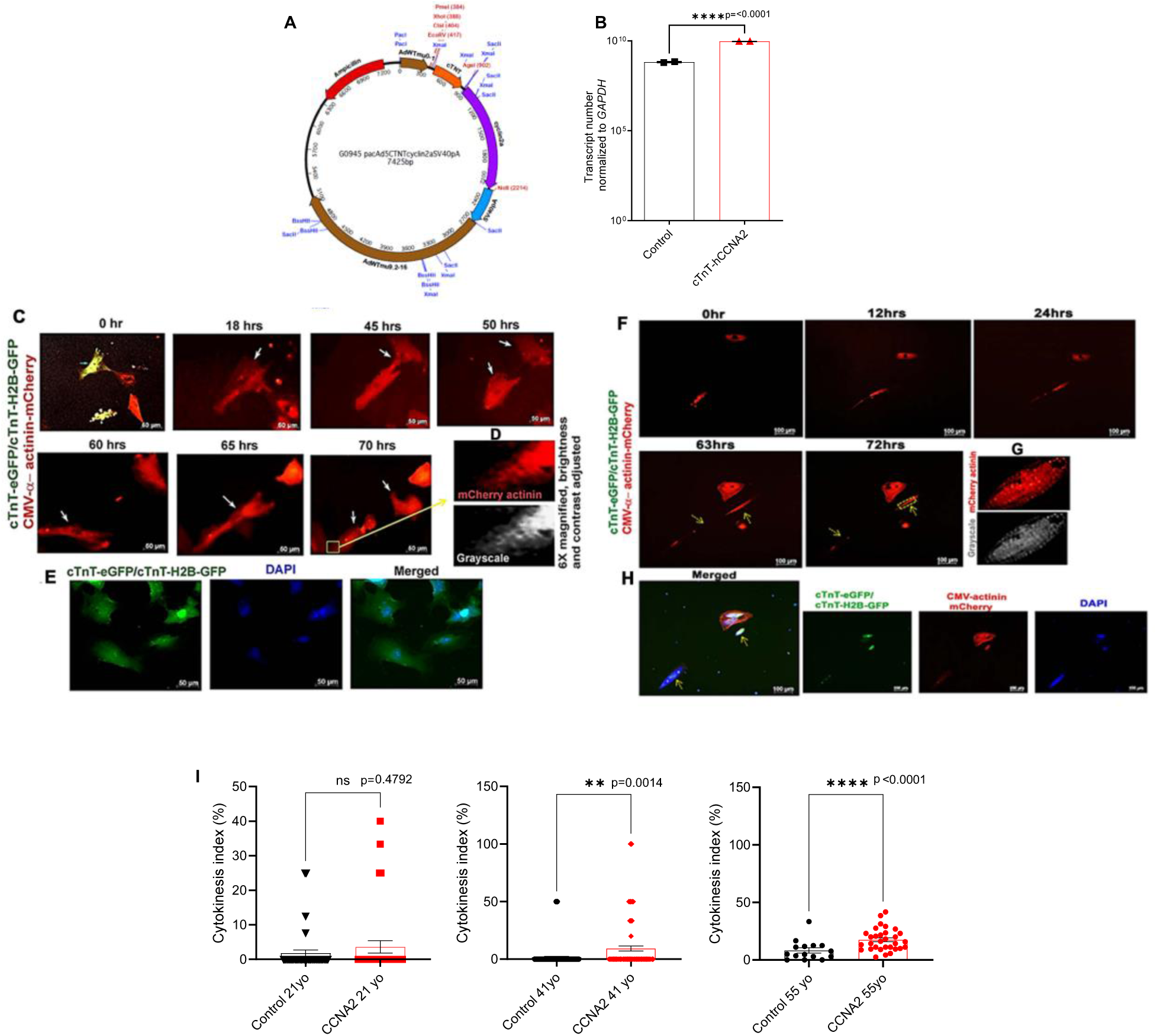
CCNA2 expression induces cytokinesis in adult human cardiomyocytes *In Vitro*. **A)** Clinical-use grade replication-deficient human adenovirus type 5 (Ad5) was used for this study. The vector was designed with cardiac Troponin T (cTnT) promoter to express human cyclin A2 (hCCNA2) specifically in cardiomyocytes. **B)** The cTnT-hCCNA2 adenovirus was used to transduce cultured adult human cardiomyocytes to induce the expression of CCNA2. Significantly higher CCNA2 expression was observed in cells transduced with cTnT-hCCNA2 compared to the control virus. **C-H)** From **Movies S1** and **S2**: still images from representative time-lapse epifluorescence microscopy of cultured cardiomyocytes isolated from adult humans **C**) 55-year-old and **F)** 41-year-old. Please note that still images of 55 and 41-years-old have scale bars of 50µm and 100µm as it is shown in the Movies S1 and S2 respectively. Cells were transduced with cTnT-hCCNA2, cTnT-H2B-GFP, cTnT-eGFP (to label cardiomyocytes), and CMV-α-actinin-m-Cherry adenoviruses prior to the beginning of the time-lapse imaging. **C)** At time 0 hr, a cell (yellow) expressing both cTnT (green) and α-actinin (red) was followed for 70 hrs via time-lapse microscopy. The green channel was closed, and cells were only followed through the red channel to avoid UV photo-toxicity. The observed human cardiomyocytes show the 1^st^ cell division at 50 hrs of imaging, and one of the daughter cells subsequently undergoes division at 70 hrs of imaging. **D** and **G)** At 70/72 hrs, a daughter cell is partially magnified with a grayscale version to demonstrate the presence of an intact sarcomere structure. **E, H)** 1 day after live imaging ended, these wells were fixed with subsequent labeling of nuclei with DAPI. The green fluorescence of the original cTnT-eGFP and cTnT-H2B-GFP transduction is visible, further confirming that these cells are cardiomyocytes. **I)** The cytokinetic events were counted in the control and test samples from the 21-, 41-, and 55-year-old subjects. A significantly higher rate of cytokinesis was observed in the test samples from the 41-year-old (*p*=0.001) and 55-year-old (*p*<0.0001) compared to the control. On the other hand, the control and test samples from the 21-year-old did not exhibit a significant difference in cytokinetic events (*p*=0.47). Data are represented as mean ± s.e.m.

### Adenoviral vector-mediated expression of CCNA2 induces cytokinesis in cultured adult human cardiomyocytes

To assess the effect of induced CCNA2 expression on cell division in adult human cardiomyocytes, we cultured adult human cardiomyocytes and studied cytokinesis *in vitro* by using live cell epifluorescence time-lapse microscopy. The adult human cardiomyocytes were plated at equal densities and transduced with cTnT-hCCNA2, cTnT-eGFP, and/or cTnT-H2B-GFP and CMV-α-act-mCherry adenoviruses (test) or with cTnT-eGFP and/or cTnT-H2B-GFP and CMV-α-act-mCherry adenoviruses (control). cTnT-eGFP and/or cTnT-H2B-GFP adenovirus was transduced in both groups to confirm the initial tracking of cardiomyocytes (green) during live cell epifluorescence microscopy. For live visualization of sarcomeric structure in cardiomyocytes, cells from both groups were co-transduced with an adenovirus containing α-act-mCherry, which we had constructed to allow for proper folding of the virally delivered α-actinin into the live cardiomyocyte sarcomere (CMV-α-act-mCherry) and was successfully used in cultured adult porcine cardiomyocytes ^22^. This strategy enabled us to confirm cardiomyocyte identity by assessing the expression of eGFP before and after cytokinesis and tracking sarcomere dynamics during live cell imaging. This approach provides a significant advantage over antibody-based identification, which is susceptible to artifacts and restricted to a single time point after cell fixation. We observed co-expression of eGFP (green) and α-actinin (red) in cultured adult cardiomyocytes **(Figure 1C** and **1H**; first panel). We performed time-lapse microscopic imaging of live cells to capture cardiomyocyte cytokinesis **(Figure 1C**, **1F**, **1H**, and **Figure S2**-a still image from **movie S1)**. The cytokinetic index of adult human cardiomyocytes was calculated by counting the cytokinetic events observed in 42 regions of interest (ROIs) **(Source Data Figure 1I)**. The cytokinetic index was significantly higher in the test samples with cTnT-CCNA2 adenovirus transduction compared to control samples in the 41- and 55-year-olds **(Figure 1I)**. In contrast, no significant change in the cytokinetic events between test and control samples was observed in the 21-year-old patient’s cardiomyocytes **(Figure 1I)**. Most remarkably, sarcomere structure was preserved in the daughter cells after cytokinesis **(Figure 1D** and **Figure 1G**; upon magnification of a daughter cell, the presence of sarcomeric structure is easily noted). The daughter cells were further identified with the expression of H2B-GFP (as they were originally also transduced with cTnT-eGFP and/or H2B-GFP) and noted to be mononuclear after they had been fixed and stained with DAPI **(Figure 1E** and **1H)**. Clusters of other cardiomyocytes with expression of eGFP could be seen adjacent to the daughter cells **(Figure 1E)**.

Next, we assessed the functional differentiation of cardiomyocytes by measuring the Ca²⁺ transients in CCNA2-expressing cardiomyocytes using live-cell calcium imaging under pacing conditions (0.5 Hz and 1 Hz; **Figure 2A**). Cardiomyocytes were maintained in long-term culture using a modified low-density protocol, followed by 3D embedding in agarose with low-serum, calcium-supplemented media. Ad5-cTnT-hCCNA2-transduced cardiomyocytes exhibited active, Ca²⁺ transients **(Figure 2B)**. These findings demonstrate that CCNA2 drives cytokinesis while preserving sarcomere integrity and calcium mobilization in adult human cardiomyocytes, suggesting functional re-differentiation.

**Figure 2.**
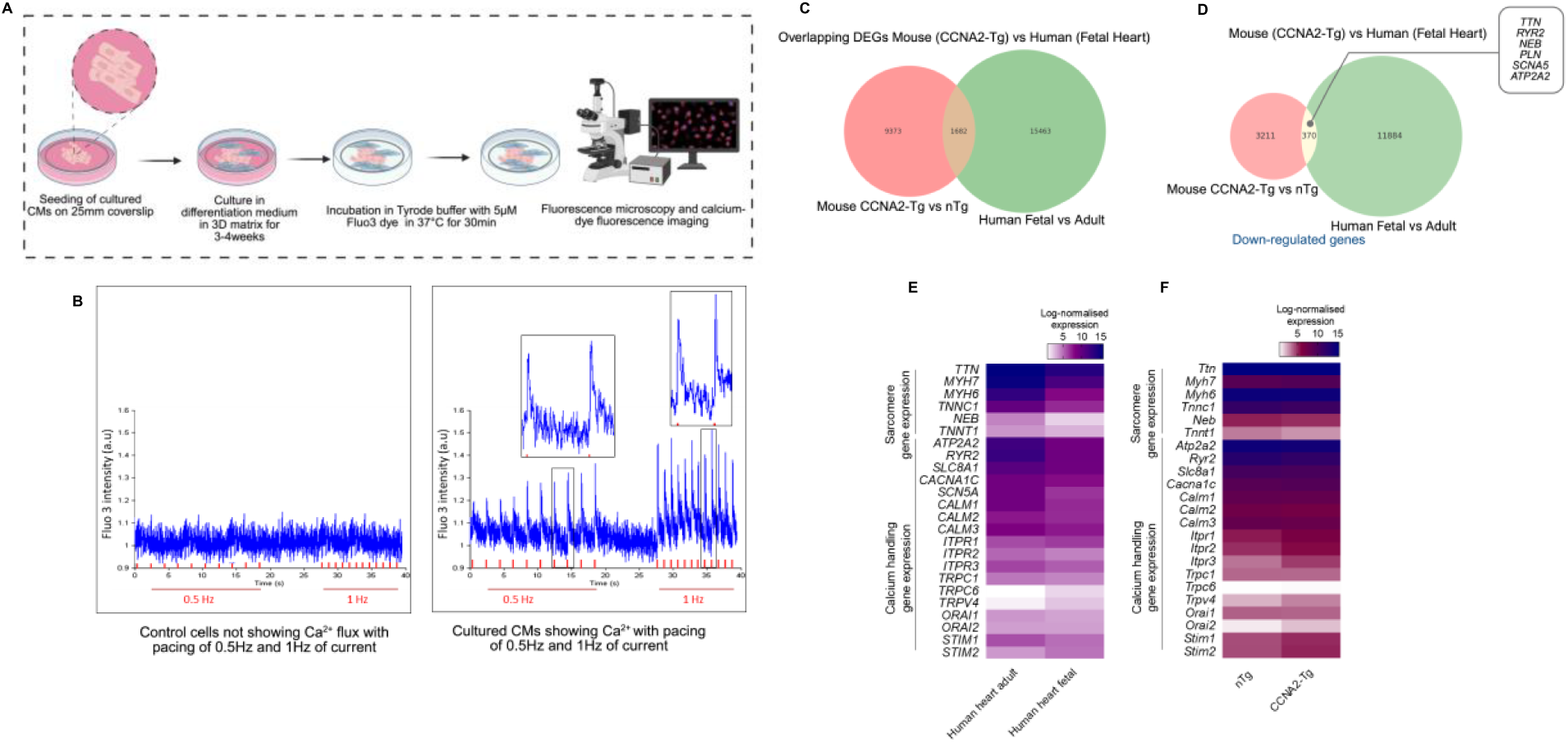
Differentiation of cultured adult human cardiomyocytes. Cultured adult human cardiomyocytes typically undergo dedifferentiation and were then subjected to 3D redifferentiation culture conditions and examined for Ca^2+^ flux. **A)** Overview of the 3D culture method of differentiation, Ca^2+^ flux measurement, and fluorescence microscopy in adult human cardiomyocytes. **B)** Representative Ca^2+^ flux graphs of re-differentiated cultured adult human cardiomyocytes. Ad5-cTnT-hCCNA2 transduced adult human cardiomyocytes treated with differentiation medium show Ca^2+^ flux with pacing of 0.5 and 1 Hz of current, while low Ca^2+^ flux was detected in the non-treated Ad5-cTnT-hCCNA2 transduced cardiomyocytes. **C)** Venn diagram showing the overlap of differentially expressed genes between CCNA2-Tg versus non-nTg mouse cardiomyocytes and human fetal versus adult hearts. **D)** Venn diagram displaying the subset of downregulated genes shared between the CCNA2-Tg mouse hearts and the fetal human heart. These genes include several associated with cardiomyocyte maturation. **E)** Heatmap depicting the expression of Ca²⁺ handling, contractility, and sarcomere-associated genes in human adult versus fetal hearts, **F)** Comparison of gene expression in cardiomyocytes from CCNA2-Tg versus nTg mice, highlighting differences in Ca²⁺ regulatory genes and contractile machinery. Positive values indicate upregulation in adult hearts relative to fetal hearts. Data represent normalized RNA-sequencing expression values. The illustration in (**A**) was created using BioRender (https://biorender.com).

### CCNA2-induced transcriptional reprogramming reveals fetal gene signatures for calcium handling and sarcomere dynamics

To quantify the developmental and reprogramming landscape, we performed ultra-deep bulk RNA-sequencing of fetal (n=3) and adult (n=4) human hearts. This approach allowed us to compare the molecular signatures of CCNA2-induced cardiomyocyte reprogramming with naturally occurring fetal-like states. Our ultra-deep sequencing approach enhanced our ability to detect low-abundance transcripts, improve quantification accuracy, and provide a more comprehensive transcriptomic profile. This comparison contextualizes how CCNA2 expression modulates gene networks associated with dedifferentiation, proliferation, and calcium handling, highlighting its role in cardiac plasticity and regeneration. Notably, there were overlapping differentially expressed genes (DEGs) between adult CCNA2-Tg mouse cardiomyocytes and fetal human hearts with downregulated genes in both that were associated with cardiomyocyte maturation **(Figure 2C** and **2D)**. Furthermore, we found that calcium handling and contractility-related genes, as well as sarcomere assembly genes such as *RYR2, ATP2A2, SLC8A1, CALM1, CALM2, NEB, TTN*, and *TNNC1,* were differentially upregulated in adult human hearts compared to fetal human hearts **(Figure 2E)**.

Additionally, we performed bulk RNA-sequencing of mouse cardiomyocytes with constitutive CCNA2 expression in cardiomyocytes (CCNA-Tg mice) compared to non-Tg mice (n=6 in total). Key calcium handling genes, including *Ryr2, Atp2a2, Slc8a1,* and sarcomere assembly genes, as well as *Neb, Ttn,* and *Tnnc1,* were significantly downregulated in CCNA2-Tg cardiomyocytes. In contrast, genes involved in intracellular calcium release and store-operated calcium entry, such as *Itpr1, Orai2,* and *Stim2*, were upregulated in CCNA2-Tg mouse hearts, further aligning with the fetal gene expression profile **(Figure 2F** and **S3A-C)**. This suggests that *CCNA2* may regulate sarcomere assembly genes during cell division, thereby facilitating cardiomyocyte proliferation ^23^. Furthermore, *CCNA2* appears to influence calcium mobilization post-cytokinesis, suggesting a broader role beyond cell cycle regulation, potentially impacting cardiomyocyte maturation and function.

Live-cell imaging of CCNA2-expression in adult human cardiomyocytes undergoing cytokinesis revealed that daughter cells retain sarcomeric integrity post-division. Despite RNA-sequencing data showing a significant downregulation of thin filament genes (*Neb, Tnn, Tnnc1*), structural sarcomere genes such as *Tnnt2* and *Actn2* remained stable **(Figure S3A)**, suggesting that sarcomere remodeling may occur transiently to accommodate cell division without leading to complete dedifferentiation.

Interestingly, transcriptomic profiling revealed upregulation of epithelial-to-mesenchymal (EMT) genes, including *Zeb1, Vim, Twist1,* and *Snai1*, which were highly expressed in embryonic (E10.5), early postnatal (P2) mouse hearts, downregulated in adult cardiomyocytes, and reactivated in CCNA2-Tg adult cardiomyocytes **(Figure S3D)**. In human fetal hearts, among EMT-associated genes, *ZEB1* was consistently upregulated in both human fetal and CCNA2-Tg hearts **(Figure S3E)**, supporting transient mesenchymal activation observed during neonatal heart regeneration ^24^. These findings suggest that CCNA2 overexpression correlates with a partial, developmentally relevant mesenchymal gene program.

We next assessed whether CCNA2 reinstates pro-proliferative gene expression along the cardiac developmental trajectory. Comparative bulk RNA sequencing analysis of E10.5, P2, and adult CCNA2-Tg and nTg cardiomyocytes revealed consistent upregulation of canonical cell cycle regulators, including *Mki67, Aurkb*, and *Cdk1,* in CCNA2-Tg cardiomyocytes relative to adult nTg controls, although their expression did not reach the embryonic and early postnatal expression levels **(Figure S4A–C)**. Despite these modest fold-changes, Cohen’s d effect size analysis revealed moderate to large positive effects for several key proliferative genes (*Mki67, Kif23, Anln,* and others) when comparing CCNA2-Tg to nTg cardiomyocytes **(Figure S4D)**, indicating that CCNA2 induces biologically meaningful transcriptional reactivation of cell cycle programs. In contrast, when comparing CCNA2-Tg cardiomyocytes to embryonic and early postnatal (P2), Cohen’s d values were negative, reflecting a high baseline proliferation in these developmental periods, and indicating that CCNA2 overexpression does not fully restore fetal gene expression levels. Notably, such partial reactivation of fetal and proliferative transcriptional programs can have significant biological effects, as demonstrated in our prior functional studies ^13,19–22^.

To validate these transcriptomic findings, RT-qPCR analysis relative to early postnatal P2 cardiomyocytes confirmed that CCNA2-Tg adult cardiomyocytes partially re-express key cell cycle and cytokinesis genes, including *Mki67, Aurka, Anln,* and *Kif23*, compared to adult nTg hearts. Notably, *Ryr2* expression was reduced in both P2 and CCNA2-Tg cardiomyocytes, reflecting a transcriptional shift toward a less mature state **(Figure S4E** and **S4F)**. These results indicate that CCNA2 reactivates proliferation without a full reversion to fetal transcriptional patterns.

Taken together, these findings suggest that the partial reprogramming induced by CCNA2 reflects a controlled activation of developmental programs, consistent with regenerative responses observed in neonatal hearts, and aligns with prior evidence that modest increases in cell cycle gene expression can enhance cardiac regeneration capacity ^13,19–22^.

### Single-nucleus transcriptomic profiling reveals CCNA2-driven reprogramming in selective cardiomyocytes, supported by integrative bulk RNA-sequencing analysis

To investigate CCNA2-induced cardiomyocyte cytokinesis in an unbiased manner, we performed snRNA-seq on hearts isolated from control non-transgenic (nTg) and *CCNA2* constitutively expressing transgenic (CCNA2-Tg) adult mice (n=8 in total). Both male and female mice aged 8-12 weeks were included. This approach enabled the identification of differentially expressed genes across cardiac clusters in both nTg and CCNA2-Tg mice in the context of CCNA2 expression **(Figure 3A**, **3B)**. Notably, snRNA-seq also captured underrepresented cell types often lost during enzymatic digestion in single-cell RNA-seq, thereby providing a comprehensive view of cardiac cellular heterogeneity ^23,25^.

**Figure 3.**
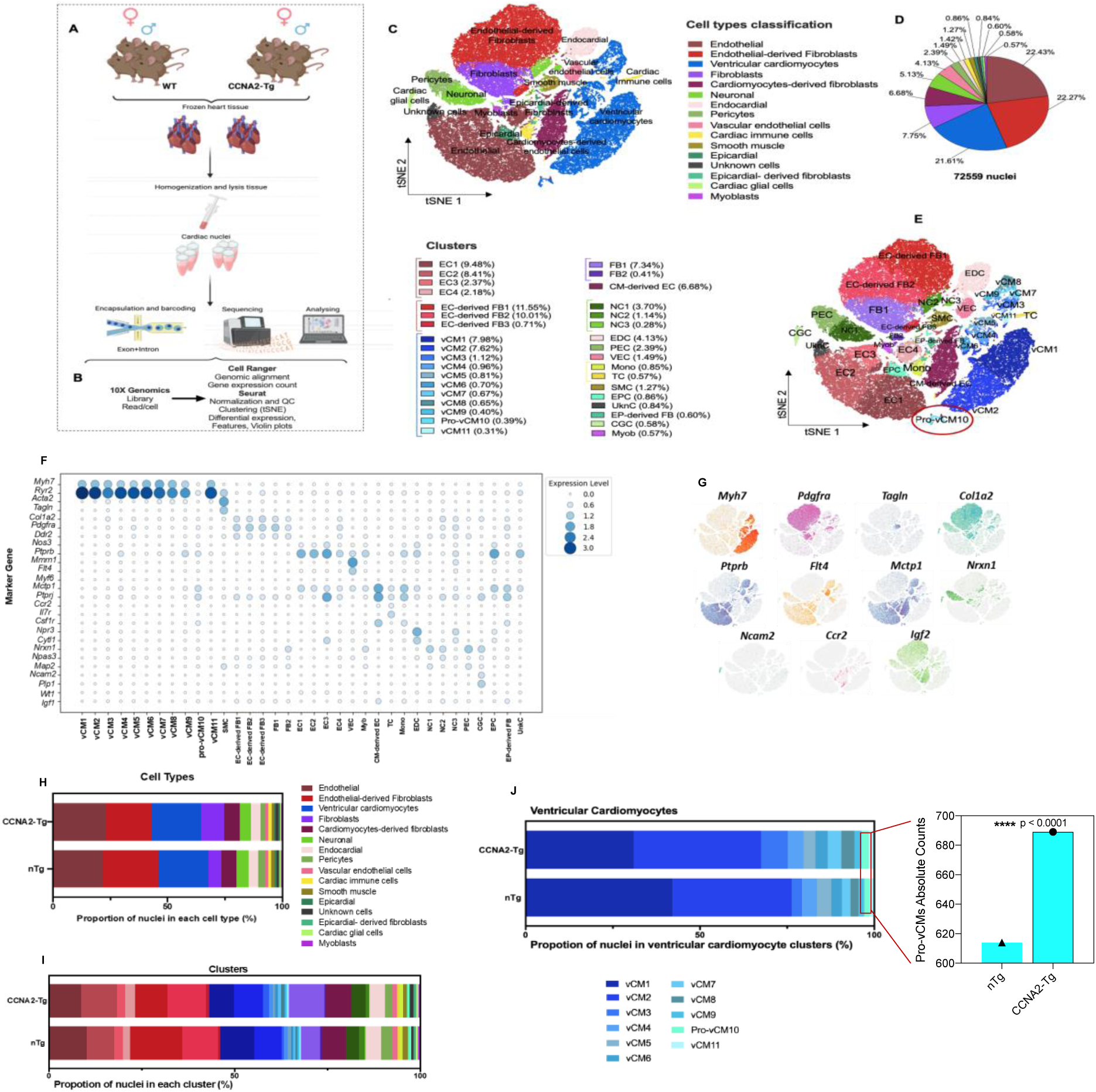
Single-nucleus RNA sequencing identified major cell types in CCNA2-Tg mice. **A)** Schematic depicting the design of the snRNA-seq experiments (n=2) for each condition nTg female, CCNA2-Tg female, nTg male, and nTg male. (*n*= 8 mice in total). **B)** The analysis pipeline and single-cell barcoded library generation (10X Genomics), sequence alignment (Cell Ranger), and further analysis using R (SeuratR package). (*n*= total nuclei 72,559) **C)** t-distributed Stochastic Neighbor Embedding (t-SNE) clustering the nuclei reveals more than 16 distinct major cell types identified based on the cell-specific markers dataset provided at iRhythmics FairdomHub instance, and **D)** the pie chart showing the proportion of cells within the snRNA-seq. **E)** t-SNE of a total of 35 subclusters. **F)** Dot plot depicting the expression levels of some of the key marker genes used to classify cell types. Dot size represents the percentage of cells expressing each gene, while color intensity reflects expression levels. **G)** Feature plots highlighting the expression of selected cell-type-specific markers across the t-SNE space. **H)** Stacked bar plots representing the proportion of nuclei classified into each major cell type (top) and **I)** each identified cluster (bottom) in both CCNA2-Tg and nTg mice. **J)** Stacked bar plots displaying the proportion of nuclei in vCM clusters (left). The absolute count comparison (right) highlights a significant increase in Pro-vCM10 in CCNA2-Tg hearts relative to nTg controls (\*\*\*\**p*<0.0001, Chi-square test), supporting the role of CCNA2 in promoting cardiomyocyte proliferation. CM: Cardiomyocytes, vCM: ventricular cardiomyocytes, EC: endothelial cells, FB: fibroblasts, EC-derived FB: endothelial-derived fibroblasts, NC: neuronal, EDC: endocardial cells. PEC: pericytes, VEC: vascular endothelial cells, Mono: monocytes, TC: T-Cells, SMC: smooth muscle cells, EPC: epicardial cells, UnkC: unknown cells, EP-derived FB: epicardial-derived fibroblasts, CGC: Cardiac glial cells, Myob: myoblasts. The illustration in (**A**) was created using BioRender (https://biorender.com).

t-distributed Stochastic Neighbor Embedding (t-SNE) of combined nuclei profiles from both nTg and CCNA2-Tg mouse hearts revealed 16 major cardiac cell populations, including cardiomyocytes, endothelial cells, fibroblasts, cardiac, immune cells, and neuronal cells, distributed across 35 subclusters (**Figure 3C**-**E**). Cell type clusters were annotated based on expression of canonical marker genes and validated through differential expression analysis and comparison with a publicly available reference dataset (iRhythmics FairdomHub) ^26,27^ **(Figure 3F** and **3G)**.

We identified several common clusters between nTg and CCNA2-Tg mice **(Figure 3E** and **Figure S6)**, with differential distribution across conditions **(Figure 3H** and **3I)**. Cardiomyocyte populations were identified by the classical marker genes (*Myh7*, *Actn2,* and *Tnnc1*) in both nTg and CCNA2-Tg hearts. Cardiomyocytes that express an enhanced level of contractility genes, including the *Ryr2*, are annotated as mature cardiomyocytes ^27^.

SnRNA-seq identified 11 subclusters of the ventricular cardiomyocyte (vCM) population common to both nTg and CCNA2-Tg hearts, with distinct differences in the proportion of nuclei across subclusters. Notably, the vCM1 and vCM5 were more prevalent in nTg hearts, whereas the vCM2, vCM3, vCM4, vCM6, vCM7, vCM8, vCM9, pro-vCM10, and the vCM11 were more abundant in CCNA2-Tg hearts **(Figure 3J)**.

Subclusters vCM1, vCM2, vCM4, vCM5, and vCM11 expressed cardiomyocyte maturation-related genes, particularly sarcomere development genes, calcium (Ca^2+^) handling, and cardiac contractility genes (*Ttn*, *Pln, Ryr2*, and *Pkp2, Kcnd2*, and *Tnni3k),* regulating cardiomyogenesis and function ^28^. Notably, vCM5 also expressed non-cardiac lineage markers such as *Ebf2*, implicated in brown adipocyte differentiation ^29^, and *Lsamp*, which guides neuronal connections ^30^. In contrast, vCM3 and vCM7 showed increased expression of ATP synthase genes (*Atp5b* and *Atp5g3),* while vCM4 and vCM8 co-expressed the endothelial-associated gene vCM9 was uniquely marked by *Igfbp3*, consistent with a mid-differentiation state.

Moreover, the transcriptomic profile of the Pro-vCM10 displayed a robust cell cycle signature with high expression of early-phase proliferative genes (*Top2a, Cdc20, Cdk2d, Cdk1, Prc1, Bub1, Mki67*, and *Kif23)*, and mitotic-cytokinesis drivers (*Stmn1*) ^31^, alongside adhesion complex assembly and immune markers (*Cd44,* and *C1qa*) ^32^. This transcriptional profile indicates an actively cycling, dedifferentiated cardiomyocyte population poised for regeneration. Remarkably, the proportion of Pro-vCM10 was significantly higher in CCNA2-Tg mouse hearts compared to nTg mouse hearts **(Figure 3J)**.

CCNA2 was expressed in cardiomyocytes, predominantly within the proliferative cluster identified in CCNA2-Tg hearts, which also expressed key cell cycle regulators **(Figure 4A** and **4B)**. This expression pattern remained consistent across the combined transcriptomic profiles of all subclusters **(Figure S7)**. The pro-vCM subcluster in CCNA2-Tg mice co-expressed proliferative markers (*Mki67* and *Kif23*) and exhibited reduced *Ryr2*, consistent with a less mature phenotype **(Figure 5A** and **5B)**. Compared to nTg cardiomyocytes, where *Ryr2* expression was higher, CCNA2-Tg cardiomyocytes expressed a transcriptional signature consistent with a dedifferentiated state that may facilitate cell cycle re-entry and regeneration ^33^.

**Figure 4.**
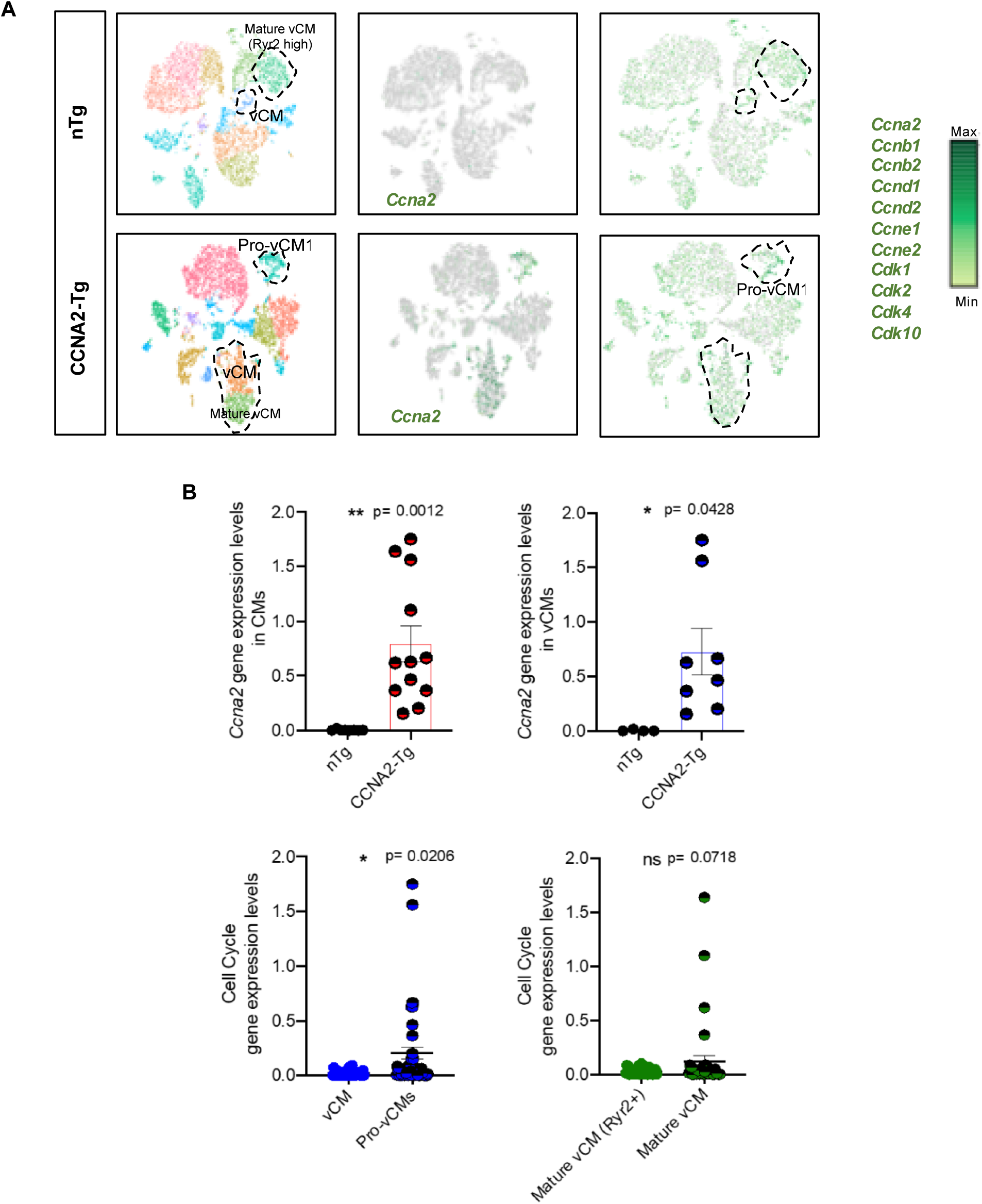
CCNA2 and cell cycle genes expression in CCNA2-Tg and nTg mice cardiomyocytes delineated by snRNA-seq. **A)** CCNA2 was expressed in the vCM, specifically in the pro-vCM subset of the cardiomyocyte population in CCNA2-Tg, and cell cycle genes projected on the t-SNE plot. **B)** Scatter plots showing the expression level of CCNA2 and cell cycle genes in cardiomyocytes of CCNA2-Tg and nTg mice. Each point represents an individual value; the mean value is represented by the horizontal line. Error bars represent s.e.m.

**Figure 5.**
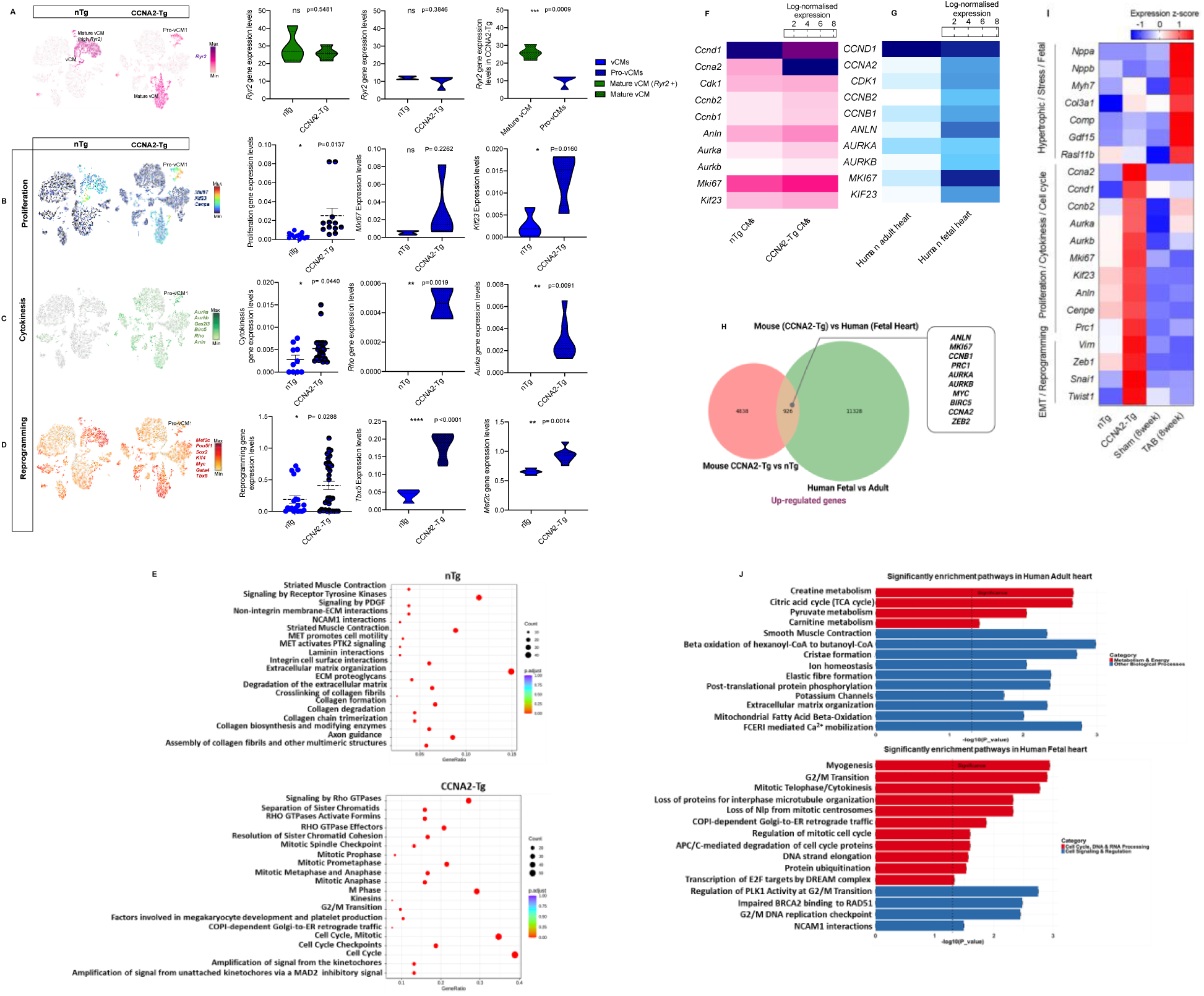
CCNA2 expression induces reprogramming, cytokinesis, and proliferation in mouse cardiomyocytes. **A)** t-SNE projection showing *Ryr2* expression across cardiomyocyte subpopulations in nTg and CCNA2-Tg mice. *Ryr2*+ cardiomyocytes correspond to a more mature ventricular cardiomyocyte (vCM) state, whereas *Ryr2* low-expression cardiomyocytes represent less mature, or proliferative cardiomyocytes. Quantification of *Ryr2* expression levels among cardiomyocyte subpopulations is shown in the accompanying violin plot, highlighting differences in *Ryr2* expression across conditions. **B)** t-SNE plots and representative scatter plots and Violin plots representing the expression of key proliferation, **C)** cytokinesis, and **D)** reprogramming gene markers in both CCNA2-Tg and nTg mice. Individual points represent median-normalized expression values for proliferation genes in Pro-vCMs. Two values were flagged as statistical outliers; these are retained to reflect biological heterogeneity within the proliferative program. **E)** Representative dot plot of the differential gene expression analysis showing the upregulation of proliferation, cytokinesis, and reprogramming genes in CCNA2-Tg versus nTg cardiomyocytes. **F)** Heatmaps of log-transformed gene expression changes in human fetal versus adult hearts, **G)** and in CCNA2-Tg versus nTg cardiomyocytes. **H)** Venn diagram showing shared upregulated genes between CCNA2-transgenic mouse hearts and human fetal hearts. The diagram displays the overlap of upregulated genes from CCNA2-Tg versus nTg mouse cardiomyocytes and human fetal versus adult heart RNA-seq datasets. These shared genes are further associated with cell cycle, cytokinesis, and cardiac development, supporting a dedifferentiation and reprogramming phenotype induced by CCNA2. **I)** Heatmap showing expression z-scores of canonical hypertrophic markers alongside genes associated with EMT and cell cycle /cytokinesis across different conditions: non-transgenic nTg, CCNA2-Tg, Sham-operated, and (8 weeks) transverse aortic banding (TAB) cardiomyocytes. The heatmap illustrates distinct gene expression patterns differentiating CCNA2-induced regenerative reprogramming from hypertrophic remodeling in pathological pressure overload. **J)** Pathway enrichment analysis showing significantly enriched pathways in the human adult (top) and fetal (bottom) heart. In the adult heart, pathways related to metabolism and energy are enriched, based on 4,348 downregulated genes and 3,713 upregulated genes (P<0.05). In the fetal heart, cell cycle, DNA/RNA processing, and signaling pathways are enriched (P<0.05).

Bulk RNA-sequencing further supported this profile, revealing coordinated downregulation of sarcomere, contractility genes, and upregulation of proliferative programs in CCNA2-Tg cardiomyocytes (**Figure S3A-C, Figure S4A-D**), aligning with a dedifferentiated state, which facilitates cell cycle re-entry and cytokinesis. Consistently, *Ryr2* was significantly downregulated in CCNA2-Tg cardiomyocytes compared to nTg controls, with expression levels approximating those observed in fetal hearts **(Figure S8A** and **Figure 2E)**. This pattern was validated by the quantitative PCR, where the *Ryr2* gene was downregulated in CCNA2-Tg cardiomyocytes and P2 hearts compared to adult nTg cardiomyocytes **(Figure S8B)**. Together, this transcriptional ‘immaturity’ may enable the proliferative potential necessary for myocardial regeneration.

To further examine CCNA2-induced cytokinesis events, we investigated whether distinct cardiomyocyte subclusters expressed genes associated with cytokinesis and/or reprogramming in CCNA2-Tg mice. Intriguingly, the pro-vCM population demonstrated upregulation of core cytokinesis regulators such as *Aurkb, Anln, Sept7,* and *Rho,* among others, with *Rho* and *Aurka* significantly elevated in CCNA2-Tg compared to nTg mice **(Figure 5C)**. This pro-vCM subcluster also showed increased expression of reprogramming genes, including *Pou5f1, Klf4, Gata4, Myc, Sox2,* and *Tbx5*, with *Pou5f1* and *Tbx5* notably increased in CCNA2-Tg hearts compared to nTg hearts **(Figure 5D)**. These findings were consistent with bulk RNA-sequencing analysis, which revealed increased expression of cytokinesis- and reprogramming-associated genes in CCNA2-Tg cardiomyocytes compared to nTg cardiomyocytes **(Figure S3D** and **5F)**.

Pathway enrichment analysis of CCNA2-Tg cardiomyocytes shows significant activation of cell cycle, cytokinesis, and mitotic regulation pathways compared to nTg controls. Notably, Rho GTPase signaling, spindle checkpoint regulation, and cytokinesis pathways were highly enriched **(Figure 5E)**, indicating increased proliferation and cytoskeletal remodeling in CCNA2-Tg hearts. In contrast, nTg cardiomyocytes showed enrichment in extracellular matrix organization, collagen biosynthesis, and muscle contraction pathways, reflecting a more differentiated state. These findings indicate that CCNA2 enhances proliferation and cellular plasticity, promoting a regenerative phenotype in cardiomyocytes.

Our bulk RNA-sequencing data from mouse nTg and CCNA2-Tg cardiomyocytes yielded comparable findings. We observed upregulation of *Ccnb1* and *Ccnb2* in CCNA2-Tg cardiomyocytes compared to nTg cardiomyocytes, along with slight increases in the expression of *Cdk1* in CCNA2-Tg cardiomyocytes **(Figure 5F)**. *Cdk1* is the sole essential cyclin-dependent kinase (*CDK*), when complexed to Cyclin A2 promotes entry into mitosis, whereas Cyclin A2/Cdk2 is critical for the G1/S transition ^13,34,35^. Key cytokinesis and proliferation genes, including *Aurka, Aurkb, Anln, Mki67,* and *Kif23,* were upregulated in CCNA2-Tg cardiomyocytes, reinforcing the role of CCNA2 in cell cycle reactivation **(Figure 5F)**. Adult human hearts exhibited downregulation of proliferative and cytokinesis-related genes **(Figure 5G** and **5H)**, whereas CCNA2-Tg cardiomyocytes retained a transcriptional profile favoring cell cycle progression and cytokinesis.

Moreover, we independently confirmed the transcriptional patterns observed in snRNA-seq, we performed RT-qPCR for key proliferation, cytokinesis genes (*Ccna2, Mki67, Kif23, Anln, Aurka)*. CCNA2-Tg cardiomyocytes exhibited elevated expression relative to nTg adults, consistent with the transcriptomics data **(Figure S8C** and **S8D)**.

To further delineate CCNA2-driven transcriptional reprogramming from pathological remodeling, we performed a comparative transcriptomic analysis using publicly available bulk RNA-sequencing data from a mouse model of pressure overload-induced hypertrophy (TAB; GSE138299). While CCNA2-Tg cardiomyocytes displayed robust induction of genes involved in cell cycle progression, mitosis, cytokinesis, and mesenchymal-like programming (e.g., *Ccna2, Ccnb2, Aurkb, Anln, Prc1, Cenpe, Mki67, Twist1, Snai1, Zeb1,* and *Vim*), hypertrophic cardiomyocytes showed selective upregulation of classical hypertrophy-associated and stress-response genes, including *Nppa, Myh7, Gdf15*, *Rasl11b,* and *Comp* **(Figure 5I)**, consistent with previous studies ^36,37^. Notably, although some cell cycle genes may be modestly increased in hypertrophy, key mitotic spindle checkpoint regulators that were strongly activated in CCNA2-Tg hearts remained low or unchanged in hypertrophic samples **(Figure S4G)**, suggesting that hypertrophy does not fully engage the cellular machinery required for successful cell division. Together, these data reinforce a key distinction between regenerative and hypertrophic responses at the transcriptomic level. These transcriptomic differences are consistent with prior functional studies demonstrating that CCNA2 gene therapy reduces fibrosis and improves cardiac function in a large-animal myocardial infarction model without incurring an increase in cardiomyocyte size ^22^, supporting the regenerative, rather than pathological, nature of CCNA2-induced remodeling.

Differential expression analysis further revealed that adult human hearts exhibit a greater number of downregulated genes compared to the fetal heart **(Figure 5J)**, consistent with widespread transcriptional silencing during cardiomyocyte maturation. In contrast, CCNA2-Tg cardiomyocytes displayed a higher number of upregulated genes relative to nTg controls **(Figure S8E)**, indicating broad transcriptional reactivation. This inverse pattern supports the role of *CCNA2* in restoring a fetal gene expression landscape.

Pathway enrichment analysis of these differentially expressed genes confirmed upregulation of metabolism, Ca^2+^ mobilization, and ion homeostasis pathways in adult human heart, and downregulation of pathways linked to mitotic progression and developmental gene programs **(Figure 5J)**. These findings support the hypothesis that CCNA2 reactivates a fetal-like transcriptional program in adult cardiomyocytes, promoting cell cycle re-entry without complete dedifferentiation.

Furthermore, pathway enrichment and gene set enrichment analysis (GSEA) revealed activation of key signaling pathways associated with cardiomyocyte plasticity, proliferation, and reprogramming in CCNA2-Tg cardiomyocytes. Specifically, the NOTCH signaling pathway and its downstream effectors (HES/HEY) were upregulated, reinforcing CCNA2’s role in progenitor-like gene activation and cardiac reprogramming, in contrast to enrichment of fibrosis, cytoskeletal remodeling, and calcineurin-NFAT signaling in TAB hearts, as previously described in transcriptomic studies of cardiac hypertrophy ^36,37^. Additionally, developmental pathways such as TGF-beta regulation and Wnt interactions suggest a shift toward a more plastic, regenerative state. Increased signaling through MAPK, PDGF, and Jak-STAT pathways further supports enhanced proliferation and cytoskeletal remodeling. Furthermore, upregulation of focal adhesion, integrin interactions, and actin cytoskeleton regulation pathways indicate structural adaptations that facilitate cardiomyocyte dedifferentiation and division. Conversely, mitochondrial metabolism and oxidative phosphorylation, hallmarks of mature cardiomyocyte function, were significantly downregulated, aligning with a metabolic reversion to a fetal-like, proliferative phenotype in CCNA2-Tg cardiomyocytes **(Figure S8F** and **S8G)**.

In summary, our data highlight roles for *CCNA2* to enhance reprogramming and dedifferentiation, which ultimately elicits cardiomyocyte cytokinesis **(Figure 6)**. These results provide a compelling pathway forward for the clinical development of cardiac regenerative therapy based on the manipulation of CCNA2 expression in cardiomyocytes.

**Figure 6.**
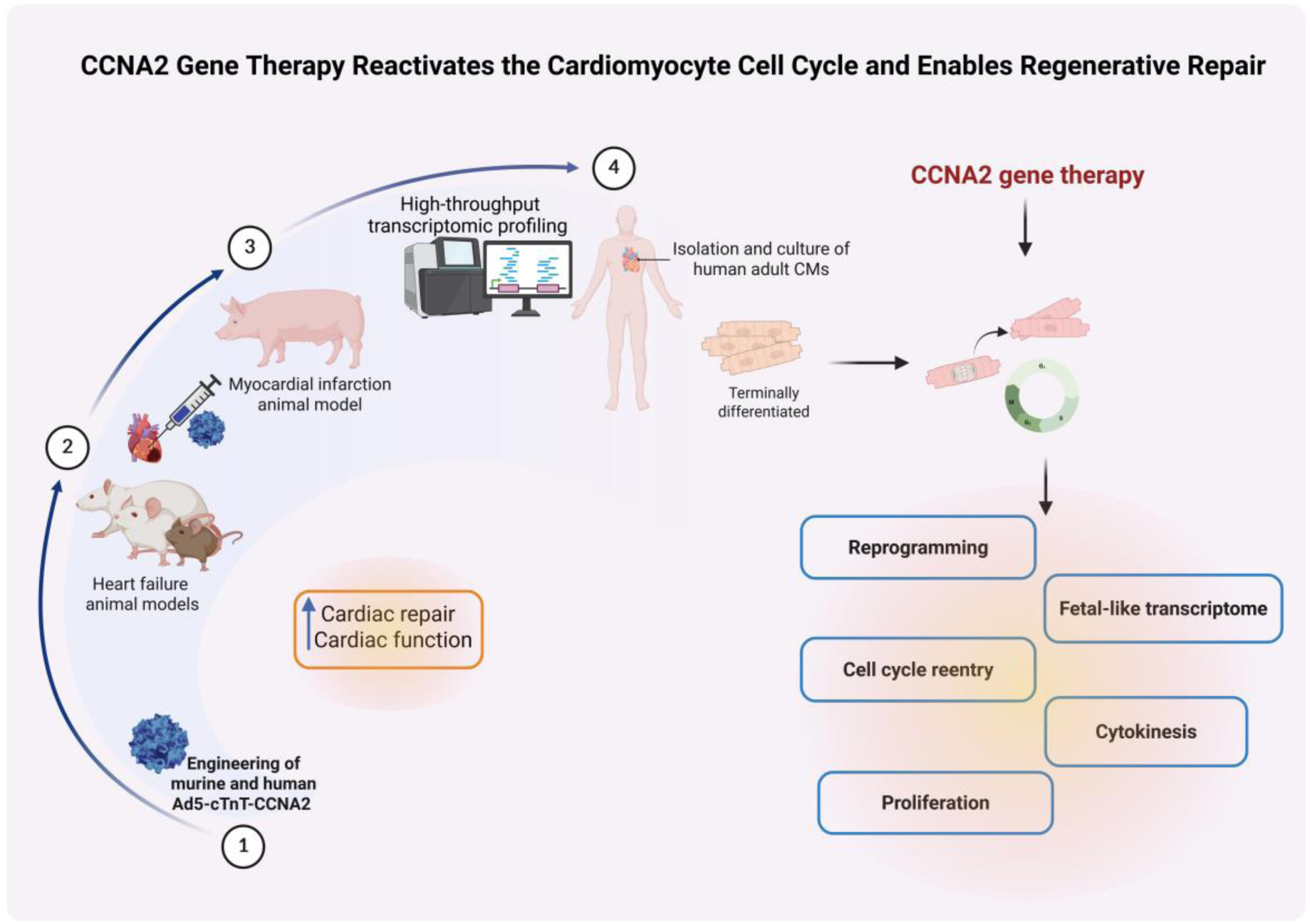
Translational roadmap and mechanistic model of CCNA2 gene therapy for cardiac regeneration. Building upon our previous *in vivo* studies in rodent and porcine myocardial infarction models, which demonstrated functional recovery via CCNA2 gene therapy (1-3) ^13,19–22^, this study provides direct evidence of complete cytokinesis in cultured human adult cardiomyocytes. Complementary transcriptomic analyses from fetal and adult human hearts, as well as CCNA2 transgenic models, reveal reactivation of reprogramming, fetal-like, and proliferative gene programs. The schematic illustrates the translational trajectory from preclinical to mechanistic insights and highlights CCNA2 as a driver of cardiomyocyte cell cycle reentry, cytokinesis, and regenerative gene expression. The illustration was created using BioRender (https://biorender.com).

## Discussion

Epimorphic regeneration is a conserved mechanism across evolution, allowing organ-specific repair in many metazoans, including neonatal mammals ^38^. In humans, this capacity is evident in neonates, where the ventricular myocardium responds to pressure overload of congenital aortic stenosis through hyperplastic growth, whereas in older patients similar stress induces only hypertrophy. ^39–41^. These clinical observations imply that the cell cycle repertoire of human neonatal cardiomyocytes is also intact, as cardiac growth is able to occur in a similar fashion to the embryonic heart.

Strategies to induce cardiomyocyte cytokinesis in adult human hearts are of critical need in order to face the growing public health crisis of congestive heart failure. To our knowledge, most cell therapy clinical trials in heart disease have provided inconsistent evidence of improvements in left ventricular ejection fraction, with a lack of evidence that the cell types utilized in patients to date actually differentiate into functional cardiomyocytes. A more direct and physiologic strategy is to reawaken proliferation within endogenous cardiomyocytes themselves. Our study provides the first demonstration that targeted expression of human CCNA2 under the control of a cardiomyocyte-specific promoter can induce complete cytokinesis in adult human cardiomyocytes, including those derived from individuals up to 55 years of age. This is a striking finding, as human cardiomyocyte turnover is thought to diminish to negligible levels by adulthood, with cytokinesis markers virtually absent beyond age 20 ^7,8^.

Importantly, our vector design—cTnT-driven CCNA2 delivered via clinical-grade Ad5—directly addresses two key translational barriers: (i) avoiding apoptosis or checkpoint failure that occurs with indiscriminate cell-cycle activation ^42–46^, and (ii) restricting CCNA2 expression to cardiomyocytes, thereby mitigating oncogenic risk in extracardiac tissues ^47^. Together, these features position CCNA2 as a next-generation gene therapy platform for safe and tissue-specific cardiac regeneration.

Mechanistically, our findings extend beyond our prior animal studies. Human cardiomyocytes isolated from 21-, 41- and 55-year-olds who had died of non-cardiac causes were cultured utilizing adult mammalian cardiomyocyte culture methods developed by our laboratory ^22^ and modified for human primary adult cardiomyocyte culture. These were then transduced with cTnT-hCCNA2. Cardiomyocytes from the 41- and 55-year-olds in the experimental wells underwent complete cytokinesis at a significantly higher frequency than the cardiomyocytes in the control wells. Based on our previous studies of rodent cardiomyocytes, we attribute the low rate of cytokinetic events we observed in the control wells to the reactivation of endogenous *CCNA2* with the prolonged culture of cardiomyocytes. This is a phenomenon we have previously observed and quantified in rodent cardiomyocytes, and likely not correlated with the low turnover seen in human hearts as measured by 14C dating ^48^ as cytokinesis markers were not noted at all in human hearts over the age 20 by Kühn and colleagues ^7^. Similarly, the Frisén laboratory ^48^ used stereological and 14C quantification techniques to measure cardiomyocyte exchange. They found that most cardiomyocytes were never exchanged and that the substantial replacement of existing cardiomyocytes was generated in the first 10 years of life, while the second decade of life focused on DNA polyploidization.

It is noteworthy to mention that in the adult murine context, approximately 85–90% of cardiomyocytes exist in a binucleated state ^49^, indicating that the nuclear count does not precisely correspond to the cell count. The polyploidization of cardiomyocytes has been recognized as an impediment to heart regeneration ^50^. It has been reported that binucleated cardiomyocytes re-entering the cell cycle are less likely to complete cytokinesis, thereby impeding the efficient regeneration of cardiac tissue following injury ^38^. In adult mammalian hearts, the ratio of binucleated to mononucleated cardiomyocytes varies from one species to another. In rodent hearts, approximately 90% are binucleated, while in human hearts, this percentage is markedly lower, varying from 25-60% ^51^. Thus, a greater percentage of mononuclear cardiomyocytes in the human heart would enable greater efficiency of CCNA2 gene therapy to induce cardiomyocyte proliferation as a repair strategy.

Our snRNA-Seq also confirms heightened cardiomyocyte plasticity of CCNA2 transgenic mice as we note changes in gene expression involved in cell cycle progression and reprogramming. Ca^2+^ signaling is a fundamental pathway in the regulation of cell division, cardiac contractility, and remodeling ^52–54^. The increase in the cytokinetic index in the test samples from 41- and 55-year-olds shows the ability of *CCNA2* to stimulate intracellular Ca^2+^ mobilization. Of note, the transduced cTnT-hCCNA2 (test samples) cultured cardiomyocytes isolated from a 21-year-old male did not show a significant increase in cytokinetic index invoked by CCNA2 when compared to the control samples, which is consistent with reports of previous investigators ^7,48^ that cell turnover and cell division can naturally occur at younger ages in humans.

SnRNA-seq has been used to identify transcriptomic changes occurring at the single-cell level, shedding light on the precise mapping of transcriptomic alterations in a cell-type-specific manner in response to cellular stimulation or gene delivery ^55^. Employing snRNA-seq in intact tissues enables the capturing of cell-specific transcriptomics changes in a more precise manner while maintaining the integrity of tissue responses, compared to isolated *in vitro* transcriptomics analyses. Our strategy of using snRNA-seq is to identify cardiomyocyte-specific changes in an *in vivo* setting in a clinically relevant CCNA2-Tg model, which allowed us to map specific changes to a subpopulation of cardiomyocytes. This would not have been captured by analyzing the transcriptome of isolated cardiomyocytes using bulk RNA-sequencing strategies. Live-cell imaging demonstrates that CCNA2-transfected adult human cardiomyocytes successfully undergo cytokinesis, with daughter cells retaining what appear to be intact sarcomeric structures. However, transcriptomic analyses (bulk RNA-sequencing and snRNA-seq) revealed significant downregulation of sarcomere-related genes. These findings suggest a dynamic remodeling process during cytokinesis, where transcriptional downregulation of sarcomeric genes may occur transiently, while structural integrity is preserved through post-transcriptional mechanisms or protein stability. This highlights the intricate balance between cellular reprogramming and maintenance of cardiomyocyte identity during division.

Importantly, while partial activation of fetal genes can occur in both regenerative and pathological contexts, our comparative transcriptomic analysis revealed a clear divergence between CCNA2-induced cardiomyocyte reprogramming and the transcriptional landscape of pressure overload-induced hypertrophy. Unlike hypertrophic hearts, which are dominated by stress-responsive and fibrotic gene signatures ^36,37^, CCNA2-Tg cardiomyocytes selectively upregulate genes involved in mitosis and cytokinesis. This pattern reflects a regenerative phenotype more akin to fetal or neonatal cardiomyocytes, rather than pathological remodeling. Notably, mitotic spindle checkpoint regulators remain largely inactive in hypertrophic models ^56^. These distinctions are further supported by pathway enrichment analyses demonstrating that CCNA2 drives activation of cell cycle and cytokinesis pathways, while hypertrophy engages calcineurin-NFAT signaling and cytoskeletal remodeling. Together, these findings support the idea that CCNA2 expression promotes a transcriptional program that is distinct from—and mechanistically incompatible with— pathological hypertrophy, further highlighting its potential as a regenerative therapeutic strategy. The combined molecular and functional evidence supports that CCNA2 expression drives a regenerative program distinct from pathological remodeling, underscoring its promise as a therapeutic strategy for myocardial repair.

Furthermore, the genes induced in the CCNA2-Tg model appear to represent a common mechanism underlying cytokinesis in both male and female mice, consistent with observations in human male and female hearts. Given that we identified a subset of cardiomyocytes that are reprogrammed into a more proliferative state, we can envision targeting this subset in future regenerative strategies. Unlike other regenerative approaches that separately target cardiomyocyte proliferation or reprogramming, our findings suggest that *CCNA2* integrates both processes, making it a unique therapeutic candidate for cardiac regeneration. Defining the transcriptomic state of cyclin A2-expressing cardiomyocytes and their proliferative potential in greater detail can enable us to recapitulate such proliferative cardiomyocytes both *in vitro* and *in vivo* to determine whether targeting these specific subpopulations holds greater therapeutic potential than targeting the entire cardiomyocyte population. The use of computational resources can aid our comprehension of these molecular mechanisms for more precision-guided regenerative strategies for human heart disease.

Another novel insight from our study is the observation that CCNA2 induces a plastic, mesenchymal-like state including upregulation of *Zeb1*, echoing mechanisms of neonatal regeneration. This transient plasticity appears to facilitate division while preserving structural integrity—a balance rarely achieved in regenerative interventions.

While these findings are promising, several limitations warrant consideration. Freshly isolated adult human cardiomyocytes, though an ideal model, are prone to culture-related cell death and variable adherence, complicating cytokinesis studies. More rapid isolation from cadaveric hearts and improved culture methods may enhance consistency. Adenoviral vectors, though used safely in clinical gene therapies, raise concerns about immunogenicity, and we are investigating alternative delivery approaches. Additionally, in vitro systems cannot fully recapitulate the in vivo cardiac environment. Our current reliance on high-resolution epifluorescence microscopy could be complemented by quantitative sarcomere scoring or super-resolution imaging to strengthen evidence of sarcomere preservation. Despite these limitations, this study represents a critical translational step, building on our prior demonstration that CCNA2 gene therapy significantly expanded cardiomyocyte numbers in porcine hearts with functional recovery, and highlights the therapeutic potential of CCNA2-induced cytokinesis in adult human cardiomyocytes.

Taken together, our results position CCNA2 as a unique therapeutic candidate that unifies cell-cycle reentry, reprogramming, and functional preservation of adult human cardiomyocytes. Unlike cell transplantation or non-specific mitogens, CCNA2 integrates proliferative drive with lineage fidelity, thereby offering a safer and more effective strategy for cardiac regeneration. Future work defining the molecular signatures of proliferative CCNA2-positive cardiomyocytes will enable the development of precision-guided therapies, whether by direct gene transfer or by reactivation of endogenous CCNA2 through antisense approaches.

By demonstrating that even middle-aged adult human cardiomyocytes can be induced to undergo complete cytokinesis, our study challenges the long-standing concept of irreversible cardiomyocyte cell-cycle exit and lays the foundation for clinically translatable, cardiomyocyte-specific regenerative therapy for heart disease.

## Methods

### Culture of adult human cardiomyocytes and *in vitro* cytokinesis study

Cardiomyocytes from adult human (21-year-old male, 41-year-old female, and 55-year-old male) heart tissue were isolated after enzymatic digestion at Anabios, San Diego, CA, and were shipped to our laboratory within 24 hours of isolation. Adult human cardiomyocytes were cultured according to our previously published adult porcine cardiomyocyte culture technique ^22^ with slight modifications. In brief, upon arrival, cells were washed with serum-free Dulbecco’s modified Eagle’s medium (DMEM) (Gibco, USA) twice, and 10^5^ cells were seeded in 100mm untreated polystyrene plates (Corning, USA). Non-adherent cells were collected every 24 hours and centrifuged at 20xg for 2 min at room temperature. Cell pellet was washed with serum-free DMEM and seeded on new polystyrene plates in modified Cardiomyocyte Culture Media (mod CMC) ^22^ formulated by adding 13% fetal bovine serum (FBS), 2.5% horse serum, 1X nonessential amino acid, 1mM sodium pyruvate, penicillin (100 U/ml), streptomycin (100 mg/ml), and fungizone (0.5 mg/ml) to DMEM/F12 (50:50). Cells were washed every day with serum-free DMEM, re-seeded in new polystyrene plates and cultured for 3 days. On day 4, they were seeded in glass-bottom 24-well tissue culture plates for 20 days, with the medium changed every 4th day. The wells with cardiomyocytes exhibiting adhesion and spreading were selected. Cells in these wells were trypsinized and counted, and 10^3^ cells per well were seeded in new glass-bottom tissue culture plates. After 2 days of culture, cells were divided into two groups (test and control) and were transduced with adenoviruses. The test group was transduced with cTnT-hCCNA2 along with cTnT-eGFP and/or cTnT-H2B-GFP and CMV-α-act-mCherry adenoviruses, while the control group was transduced with only cTnT-eGFP and/or cTnT-H2B-GFP and CMV-α-act-mCherry adenoviruses. MOI of adenoviruses was adjusted to 180 in each well of the test (with cTnT-hCCNA2; MOI 100, CMV-α-act-mCherry; MOI 40, and cTnT-eGFP or cTnT-H2B-GFP; MOI 40) and the control (cTnT-eGFP; MOI 140 and CMV-α-act-mCherry; MOI 40) group. After 48 hours of incubation, transduction was confirmed by observing the desired fluorescence in live cell imaging with Zeiss AxioVision Observer Z1 inverted microscope (Carl Zeiss). We also tested the differentiation potential (Ca^2+^ flux) of daughter cardiomyocytes after cytokinesis *in vitro* by culturing them in an agarose-based semi-solid medium for 2 weeks. For calcium imaging, they were seeded on a glass coverslip and grown in low serum (5% FBS without horse serum) and 0.5 mM calcium chloride for 10 days in a 3D matrix (agarose). For control cells (also daughter cells post cytokinesis with CCNA2), 0.5 mM calcium chloride was not added. Cells were incubated in Tyrode buffer containing 5µM Fluo3 dye for 30 minutes. Cells were washed with Tyrode buffer. Cardiomyocytes were subjected to Ca^2+^ flux imaging with pacing of 0.5 Hz and 1 Hz current. The flux was graphed as time (s) on the X-axis and Fluo3 intensity (Ca^2+^) on the Y-axis.

### Time-lapse microscopy

To capture cell division events in cardiomyocytes *in vitro*, we carried out live cell epifluorescence time-lapse microscopy as described in ^22^. In brief, Zeiss AxioVision Observer Z1 (Carl Zeiss, Thornwood, NY, USA) inverted epifluorescence microscope in a humidified chamber in the presence of 5% CO_2_ at 37°C was used to carry out time-lapse microscopy. Multiple random points with cells expressing eGFP (green) and mCherry (red) were selected in the test and control groups. The positions were marked with the “position-list” tool in the AxioVision microscopy software (AxioVision Release 4.7, Carl Zeiss). After the first cycle of imaging, only the channel for Texas red was used (for detection of mCherry) to acquire images for 72 hours. The fluorescein isothiocyanate (green) channel of the microscope was kept closed during the time-lapse imaging to avoid cell death from exposure to ultraviolet rays in this channel. Images were taken at intervals of 30 min. The 10X objective was used for all time-lapse imaging. Time-lapse movies were generated after the end of each experiment and exported as MOV files. The time-lapse movies were analyzed, and cells that underwent successful cytokinesis were enumerated in each group. The percentage (%) of cytokinesis events was calculated for each position, and the graph was plotted.

### Cell fixation and nuclear staining

After time-lapse microscopy, cells in the glass-bottom plate were fixed with 4% paraformaldehyde at room temperature for 20 min and were stored at 4°C. For nuclear staining, cells were washed with 1x PBS then permeabilized with 0.5% Triton X-100 solution for 20 min at room temperature. Cells were washed three times with 1x PBS and incubated in DAPI solution (2.5 µg/ml) for 5 min. Cells were washed twice with 1x PBS, and imaging was carried out using a Zeiss AxioVision Observer Z1 inverted epifluorescence microscope.

### Quantitative RT-PCR and Conventional PCR

Quantitative PCR was performed in human adult cultured cardiomyocytes transduced with cTnT-hCCNA2 (test) or cTnT-eGFP (control) adenoviruses with similar MOI (MOI 100). Reactions were run on a StepOnePlus™ Real-Time PCR System (Applied Biosystems) using SYBR Green PCR Master Mix (Applied Biosystems, Cat. #4309155). The PCR protocol consisted of: 95 °C for 2 min; 39 cycles of 95 °C for 15 s and 60 °C for 1 min; followed by melt-curve analysis. Gene expression was quantified using the comparative Ct (ΔΔCt) method ^22,57^ with normalization to *GAPDH*. A Ct value of 39 was assigned to reactions with a single detectable transcript. Fold change was calculated as 2^(39−ΔΔCt)^, consistent with this detection threshold approach.

For mouse experiments, total RNA was isolated from CCNA2 transgenic (CCNA2-Tg), adult non-transgenic (nTg) cardiomyocytes, and nTg postnatal day 2 (P2) hearts. RNA concentration was determined using a Qubit 3.0 Fluorometer (Thermo Fisher Scientific). First-strand cDNA was synthesized from total RNA using the High-Capacity cDNA Reverse Transcription Kit (Applied Biosystems, Cat. #4368814) according to the manufacturer’s instructions. Conventional PCR was performed with mouse-specific primers for proliferation, cytokinesis, and reprogramming markers **(Table S1)**, with *Gapdh* as the internal control. The PCR cycling conditions included: 95 °C for 10 min; 40 cycles of 95 °C for 30 s, 60 °C for 30 s, and 72 °C for 10 min; final hold at 4 °C. For quantitative PCR, 10 ng cDNA was used per reaction with SYBR Green PCR Master Mix on the StepOnePlus™ Real-Time PCR System. Data were normalized to endogenous control genes and analyzed by the comparative Ct method as above ^22,57^. Experiments were performed in triplicate.

### Animal

Cyclin A2 transgenic mice, adult males, and females 8-12 weeks old (n=4), were maintained in a B6CBA background ^13, 19^. Non-transgenic (wild-type) littermates were used as controls (n=4). Animals were housed in a temperature-controlled room with a 12:12 light/dark cycle and access to water and food. All animal experiments were conducted in compliance with ARRIVE 2.0 guidelines. In selected experiments, animals received an intraperitoneal injection of heparin (1000 U/mL; Sigma, H3393-50KU) to prevent coagulation during heart isolation for cardiomyocyte isolation, and culture. Anesthesia was induced with isoflurane (Dechra) inhalation until loss of pedal reflex confirmed deep anesthesia. While under deep anesthesia, mice were euthanized by cervical dislocation, in accordance with the current AVMA guidelines, and all procedures were performed to minimize pain and distress. A complete ARRIVE Essential 10 checklist is provided in **Table S2**.

### Collection of CCNA2-Tg mouse hearts and sample preparation for single-nucleus RNA-sequencing (snRNA-seq)

Frozen heart tissues from CCNA2-Tg- and nTg adult mice, male and female, were minced and transferred into a Dounce homogenizer containing ice-cold lysis buffer (Benthos Prime Central, TX, USA). Samples were homogenized gently, and nuclei were isolated. Nuclei were ensured to be free of clumps and debris by trypan blue staining under a microscope. Nuclei were then counted, and the concentration was adjusted. 10x library concentration, insert size, and quantification were checked using Qubit, Agilent bioanalyzer, and q-PCR, respectively. For data Processing and Quality Control of 10x Sequencing of snRNA-seq, the reads were processed with 10X Genomics CellRanger software (v.3.1.0) with the default parameters for each sample. Cell Ranger employs the STAR aligner for splicing-aware mapping of reads to the genome. It utilizes a transcript annotation GTF file to categorize the reads into exonic, intronic, and intergenic regions based on whether the reads align confidently with the genome. Only confidently mapped, non-PCR duplicates with valid barcodes and UMIs were used to generate a gene-barcode matrix. This involved various steps, including alignment to a reference, collapsing of unique molecular identifiers (UMIs), UMI counting, and initial quality control procedures. The output of this process was the generation of filtered gene expression matrices that exclusively contained cellular barcodes. Subsequently, the Seurat R package (v.3.1.1) was used for data quality control and downstream processing. The filtered_feature_bc_matrix generated by Cell Ranger was used for Seurat processing. Post-sequencing quality control was performed using Seurat. Cells were excluded from downstream processing in each sample if they met any of the following criteria: fewer than 200 genes were detected per cell, genes had non-zero counts in at most three cells, total feature counts exceeded 8000 (suggesting potential multiplets), more than 50% of the feature count was attributable to mitochondrial genes, or more than 5% of the feature count was attributable to hemoglobin-related genes, and doublets from all the clusters were filtered during QC using DoubletFinder version 2.0.3 **(Figure S5)**. Raw read counts were log-normalized, cells were clustered **(Figure S6)**, and markers for each cluster were identified by using the Seurat FindMarkers and FindAllMarker functions ^58^. The cell clusters were annotated by using the iRhythmics FairdomHub instance (https://fairdomhub.org/studies/739) and prior knowledge.

### Human fetal and adult hearts and RNA sequencing

Human fetal heart RNA (n=3 pooled, Cat #636532, Takara) and human adult heart RNA (n=4 pooled, Cat #636583, Takara) were used for bulk RNA sequencing with an increased read depth (Ultra-deep sequencing), generating approximately 300 million reads per sample. This high sequencing depth enabled the detection of rare transcripts, improved quantification accuracy, and enhanced the resolution of transcript isoforms, including novel and unannotated RNAs. For library preparation, we employed a ribosomal RNA (rRNA) depletion strategy instead of poly(A) enrichment to capture both polyadenylated (polyA⁺) and non-polyadenylated (polyA⁻) RNAs. While polyA⁺ selection primarily enriches for mRNAs and some lncRNAs, rRNA depletion allows for the inclusion of a broader spectrum of regulatory RNAs, including enhancer RNAs (eRNAs), small nuclear RNAs (snRNAs), and uncharacterized lncRNAs that may lack poly(A) tails. Additionally, this approach reduces transcript bias, preserves isoform diversity, and is better suited for degraded RNA samples.

By combining rRNA depletion with ultra-deep sequencing, we achieved a comprehensive and unbiased profile of the cardiac transcriptome. The sequence reads were mapped to the human genome (GRCh38.110) using STAR ^59^ (version 2.7.9a). Using the mapped reads, the R package edgeR ^60^ (version 3.30.3) was employed to calculate counts per million (CPM) values and false discovery rate (FDR)-adjusted p-values obtained through the Benjamini–Hochberg method (Benthos Prime Central, TX, USA).

### Comparative transcriptomic analysis of CCNA2-Tg and hypertrophic (TAB) cardiomyocytes

To delineate the transcriptional distinctions between regenerative and pathological remodeling, we compared CCNA2-Tg with cardiomyocytes from the murine transverse aortic banding (TAB) model (GSE138299) ^37^. Bulk RNA-seq datasets were processed using counts per million (CPM), followed by geometric mean normalization based on validated, stably expressed housekeeping genes *(Rpl13a* and *Hprt1)*. This approach enabled a quantitative comparison of gene expression programs underlying CCNA2-driven regeneration versus stress-induced hypertrophic remodeling.

### Pathway enrichment analysis

Pathway enrichment analysis was performed using the Reactome pathway analysis tool ^61^. The differentially expressed genes among nTg and CCNA2-Tg conditions (log2FC > / < 1 and a P value < 0.05) were used for pathway analysis. Fold enrichment was calculated to indicate the gene expression observed in our list compared to the expected list. Categories with a fold enrichment greater than 1 were considered overrepresented, while those with a fold enrichment less than 1 were considered underrepresented in our experiment. Additionally, pathway enrichment analysis was conducted using BioPlanet via the EnrichR suite ^62^, following a previously described approach ^63^.

### Statistical analysis

Statistical analyses were carried out using the R package (version 3.30.3), GraphPad Prism (version 8.2.1, GraphPad), and Python (via Google Colab). Two-tailed unpaired or paired *t*-tests were used for comparison between groups with normally distributed data. Chi-square (χ^2^) tests for proportions were performed using Python libraries, including *Scipy*. The sample size (n) is reported for each analysis. Data are presented as means ± s.e.m unless otherwise indicated. Statistical significance was defined as P < 0.05. Effect sizes were computed using Cohen’s d to assess the magnitude and directionality of expression changes between groups ^64^, defined as the difference between group means divided by the pooled standard deviation. The graph was generated in Python using Google Colab. We followed standard conventions for interpreting effect size magnitude: small (∼0.2), medium (∼0.5), and large (≥0.8). In this study, |d| ≥ 1.0 was considered indicative of a strong effect.

### Study approval

This study was conducted in accordance with the Declaration of Helsinki. Adult human cardiomyocytes were obtained from Anabios (San Diego, CA), which provides de-identified patient information as defined in the Health Insurance Portability and Accountability Act (HIPAA). Anabios consent procedures utilized meet the standard guidelines concerning informed consent for research. For the mouse study, the protocol was approved by the Institutional Animal Care and Use Committee (IACUC) of Mount Sinai Hospital, and all procedures adhered to the Institutional Animal Care and Use Guidelines.

### Data availability

The processed single-nucleus data and the deep bulk RNA sequencing, including raw expression matrices and raw sequence files that support this research study’s findings, are available on the Gene Expression Omnibus GSE249433 and GSE256519, respectively. For comparative analyses, publicly available transcriptomic data from hypertrophic hearts were obtained from GSE138299 ^37^.

## Supporting information

The real-time, live-imaging movie of human adult cardiomyocytes (41-year-old Female) is described in Figure 1

The real-time, live-imaging movie of human adult cardiomyocytes (55-year-old male) is described in Figure 1

## Abbreviations and Acronyms

Actn2: Actinin Alpha 2
Atp2a2: ATPase, SERCA2
Atp5b: ATP Synthase, Mitochondrial F1 Complex, Beta Polypeptide ATP
Atp5g3: Synthase, Mitochondrial C Subunit, Gamma Polypeptide 3
Bub1: BUB1 Mitotic Checkpoint Serine/Threonine Kinase
C1qa: Complement C1q A Chain
Ca^2+^: Calcium ion
Calm1: Calmodulin 1
Calm2: Calmodulin 2
CCNA2: Cyclin A2
CCNA2-Tg: CCNA2-transgenic mice
Cdc20: Cell Division Cycle 20
Cdk1: Cyclin-Dependent Kinase 1
Cdk2: Cyclin-Dependent Kinase 2
CGC: Cardiac glial cells
CMs: Cardiomyocytes
CPM: Counts per million
CTnT: Cardiac Troponin T
CTnT-hCCNA2: Cardiac Troponin T-human cyclin A2
E1/E2: Early region 1/Early region 2
EC: Endothelial cells
EC-derived FB: Endothelial-derived fibroblasts
EDC: Endocardial cells
Efnb2: Ephrin B2
eGFP: Enhanced Green Fluorescent Protein
EPC: Epicardial cells
EP-derived FB: Epicardial derived fibroblasts
ES: Embryonic stem cells
FB: Fibroblasts
FBS: Fetal Bovine Serum
FDR: False discovery rate
Gata4: GATA Binding Protein 4
H2B: Histone 2B
Igfbp3: Insulin-Like Growth Factor Binding Protein 3
iPS: Induced pluripotent
Kcnd2: Potassium Voltage-Gated Channel Subfamily D Member 2
Kif23: Kinesin Family Member 23
Klf4: Kruppel Like Factor 4
MI: Myocardial infarction
Mki67: Marker of Proliferation Ki-67
MOI: Multiplicity of Infection
Mono: Monocytes
Myc: MYC Proto-Oncogene, BHLH Transcription Factor
Myh7: Myosin Heavy Chain 7
Myl2: Myosin Light Chain 2
Myob: Myoblasts
NC: Neuronal cells
Neb: Nebulin
nTg: Non-transgenic mice
PEC: Pericytes
Pkp2: Plakophilin 2
Pln: Phospholamban
Pou5f1: POU Class 5 Homeobox 1, Oct4
Prc1: Protein Regulator of Cytokinesis 1
Pro-vCM: Proliferative ventricular cardiomyocyte
Ryr2: Ryanodine Receptor 2
Scn5a: Sodium Voltage-Gated Channel Alpha Subunit 5
SMC: Smooth muscle cells
SnRNA-seq: Single nucleus RNA sequencing
TC: T-Cells
Tnnc1: Troponin C1
Tnni3k: Troponin I Interacting Kinase
Top2a: Topoisomerase (DNA) II Alpha
t-SNE: T-distributed Stochastic Neighbor Embedding
Ttn: Titin
UMI: Unique molecular identifier
UnkC: Unknown cells
vCM: Ventricular cardiomyocyte
VEC: Vascular endothelial cells
α-act: Alpha-actinin

## Acknowledgment

We thank Dr. Bernhard Kühn for his contribution to the cTnT-H2B-GFP utilized in this study. This work was supported by the National Institutes of Health National Heart, Lung, and Blood Institute (NIH R01 HL150345), the New York Stem Cell Science (NYSTEM C029565 and C32608GG).

## Author Contributions

E.B. wrote the original article, contributed to experiments, and analyzed and performed snRNA-seq data with input from P.M. and A.K.; S.V.M and A.K.R contributed to cytokinesis experiments and calcium measurements; P.M. contributed to processing snRNA-seq samples and initial analysis; C.D.S. assisted with the cell biology experiments and organization of the manuscript; P.E.M., A.G., N.A.G. provided cardiomyocytes through their proprietary technology. H.W.C. was responsible for conceptualization, vector design, validation, supervision, reviewing, and funding acquisition. Review & editing-E.B, H.W.C, P.M., and A.K.

