## Supplementary Information for "Cyclin A2 Induces Cytokinesis in Human Adult Cardiomyocyte and Drives Reprogramming in Mice"

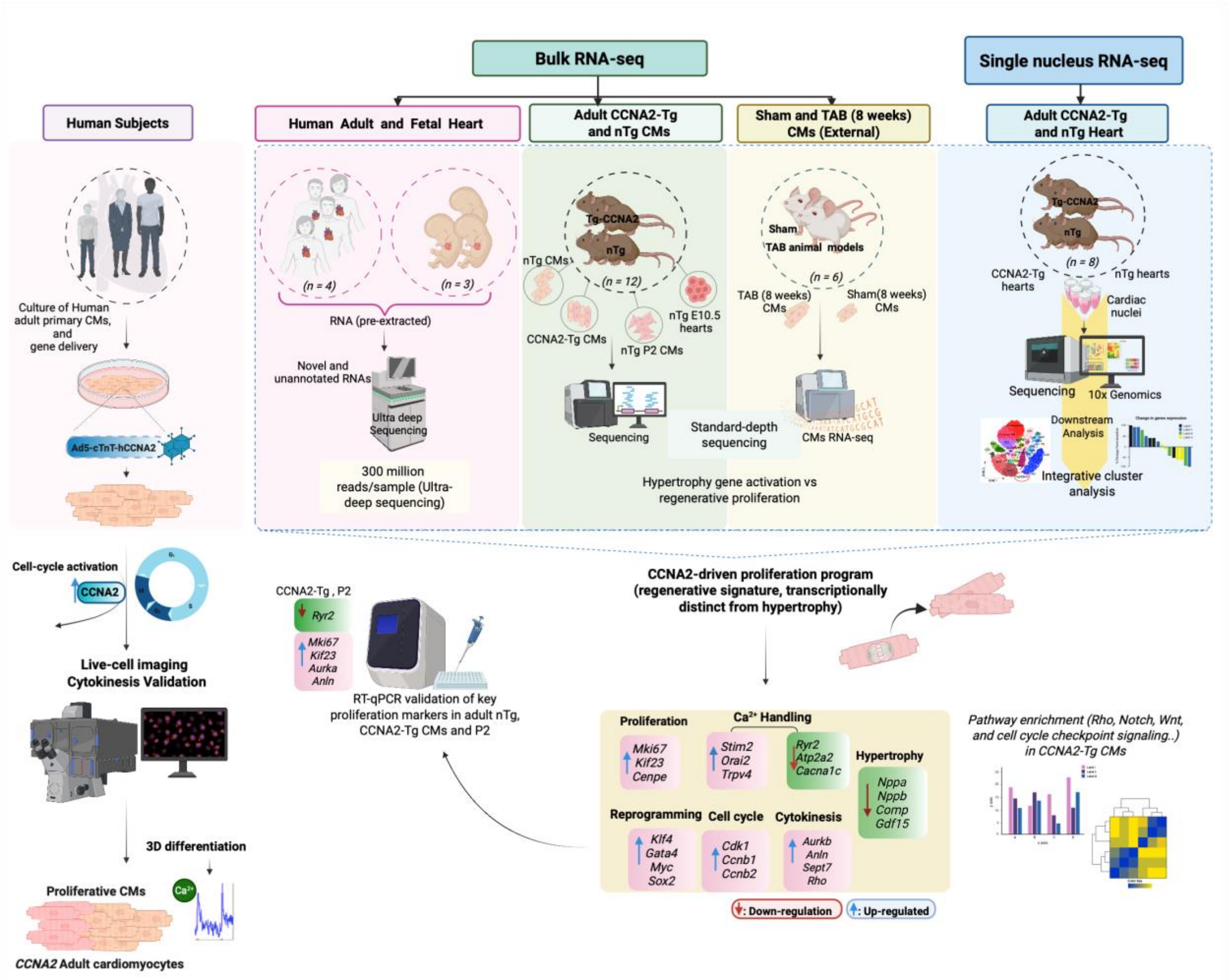


**Figure S1. Schematic overview of experimental strategy and key findings of CCNA2-driven cardiomyocyte proliferation, reprogramming, and cytokinesis validation across models.** The illustration was created using BioRender (https://biorender.com).

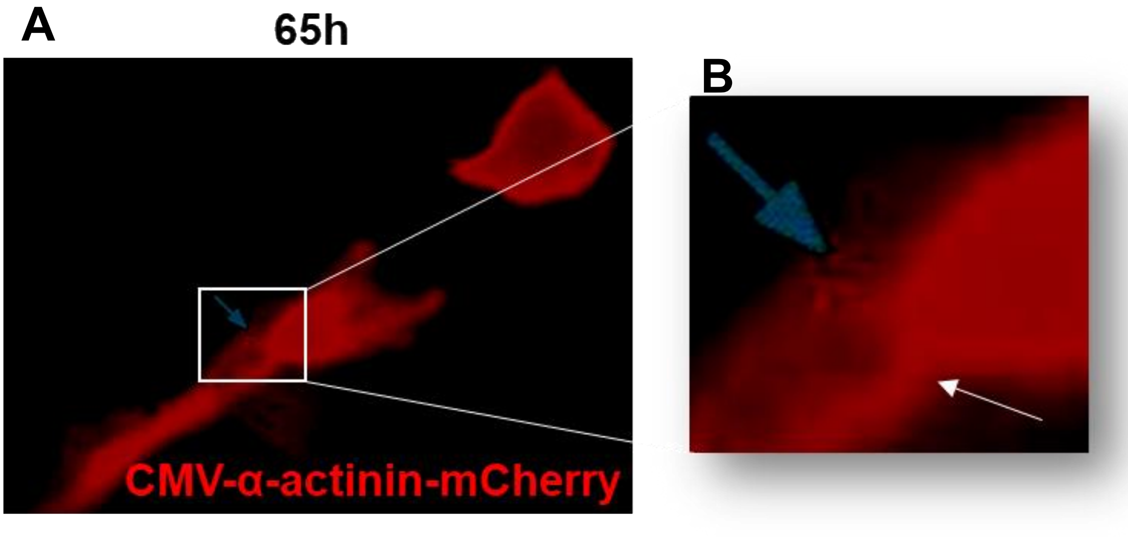


**Figure S2. CCNA2 drives cytokinesis in adult human cardiomyocytes.**
**A)** Representative still image derived from movie S1 of adult human cardiomyocytes from a 55-year-old subject and transduced with cTnT-hCCNA2, showing contractile ring formation between two dividing cardiomyocytes at 65 h post-transduction. The observation of a contractile ring highlights active cytokinesis in adult human cardiomyocytes. Please note this is not ‘fixed cell immunofluorescence’. **B)** Magnified view with arrows highlights the contractile ring.

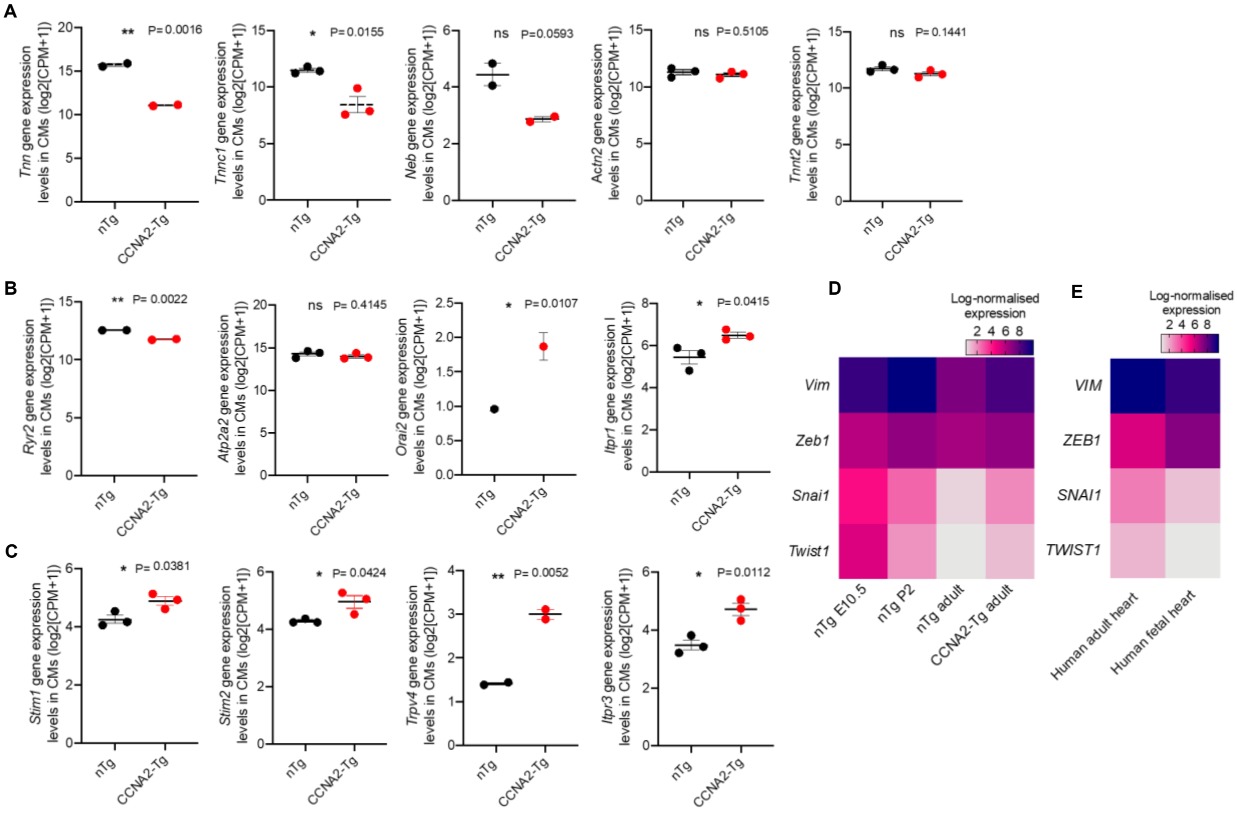


**Figure S3. Gene expression changes in CCNA2-Tg cardiomyocytes. A)** Representative scatter plots of sarcomere assembly genes *(Neb, Tnn, Tnnc1, Actn2,* and *Tnnt2),* and **B)** $Ca^{2+}$ handling genes (*Ryr2, Atp2a2, Slc8a1*, and others) in nTg and CCNA2-Tg cardiomyocytes as delineated from bulk RNA seq. Each point represents an individual value; the mean value is represented by the horizontal line; error bars represent s.e.m, with statistical significance indicated (*P*-values). “ns" denotes non-significant comparisons. **C)** Heatmap showing log-normalized expression of mesenchymal transition markers (*Vim, Zeb1, Snai1,* and *Twist1*) across embryonic (E10.5), postnatal (P2), adult nTg, and CCNA2-Tg cardiomyocytes. **D)** Heatmap of corresponding gene expression in human adult and fetal hearts. Expression values are log-transformed and normalized across samples.

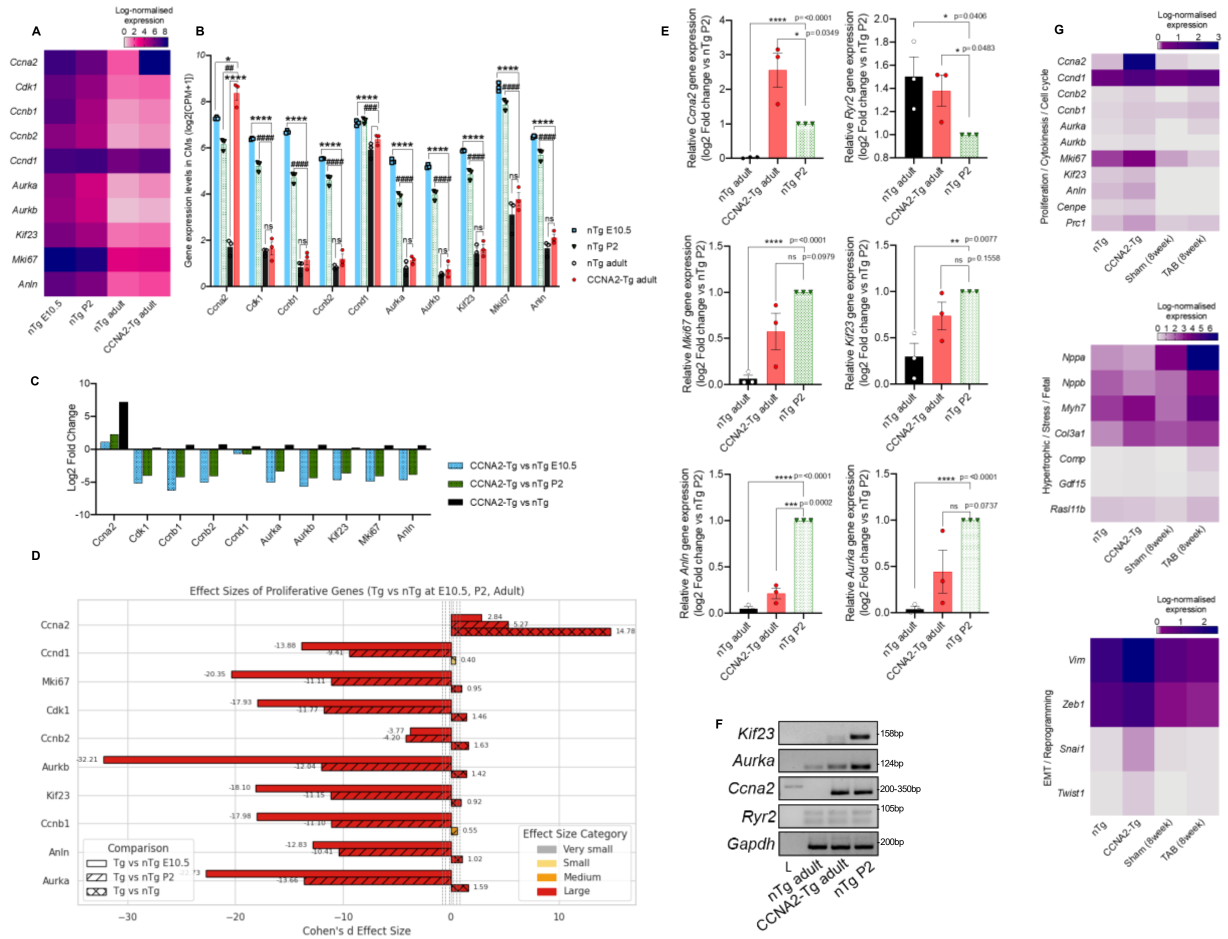

**Figure S4. CCNA2 overexpression promotes proliferation-associated gene expression in cardiomyocytes. A)** Heatmap showing expression of cell cycle/proliferation-associated genes (*Ccna2, Ccnb1, Ccnb2, Aurka, Aurkb, Cdk1, etc.*) across developmental stages (E10.5, P2, adult nTg, and adult CCNA2-Tg cardiomyocytes) from mouse bulk RNA-seq. **B)** Scatter plot representation of the same proliferation-associated genes shown in (A), expressed as $\log_2$(CPM+1). Each dot represents an individual biological replicate from bulk RNA-seq, and bars indicate mean ± s.e.m. **C)** $\text{Log}_2$ fold change in proliferation-associated gene expression in CCNA2-Tg versus nTg cardiomyocytes and across developmental stages. **D)** Effect size (Cohen's d) for proliferation-associated genes across developmental transitions (Adult CCNA2-Tg vs E10.5; Adult CCNA2-Tg vs P2; Adult CCNA2-Tg vs adult nTg). **E)** RT-qPCR validation of proliferation-associated genes in P2, adult nTg, and CCNA2-Tg cardiomyocytes. Expression values were normalized to *Gapdh* and are presented as fold change relative to nTg P2 (mean ± s.e.m.; n=3 independent experiments; ROUT outlier detection Q=1%). **F)** Representative conventional PCR electrophoresis analysis of proliferation markers (e.g., *Kif23, Aurka, Ccna2, Ryr2*) in nTg versus CCNA2-Tg cardiomyocytes, with *Gapdh* as loading control. **G)** Heatmaps showing expression canonical hypertrophic markers alongside genes associated with EMT and cell cycle /cytokinesis across different conditions: adult nTg, CCNA2-Tg, Sham-operated, and (8 weeks) transverse aortic banding (TAB) cardiomyocytes. Expression values are shown as log-normalized counts.

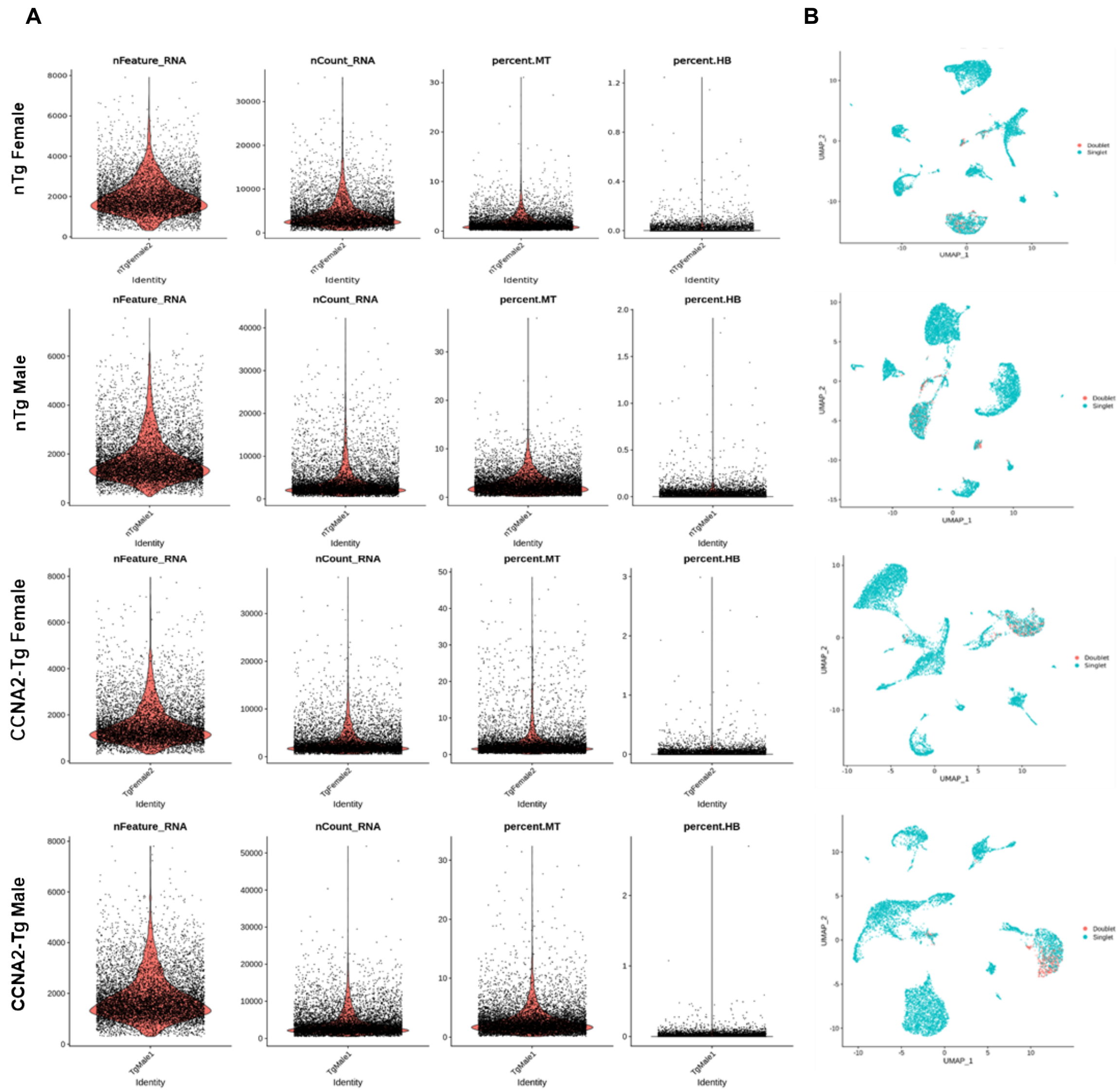


**Figure S5. Quality control (QC) filtering and visualization in SnRNA-seq. A)** "nFeature_RNA" indicates the count of gene features; "nCount_RNA" represents the count of UMIs; "percent.MT" signifies the proportion of counts originating from mitochondrial genes, and "percent.HB" signifies the proportion of counts stemming from Hemoglobin-related genes in all conditions nTg and CCNA2-Tg males and females. Each data point corresponds to a cell. **B)** The UMAP plots are generated from the top principal components within the clustered dataset. Each data point represents an individual cell and is color-coded based on whether it is categorized as a singlet or doublet.

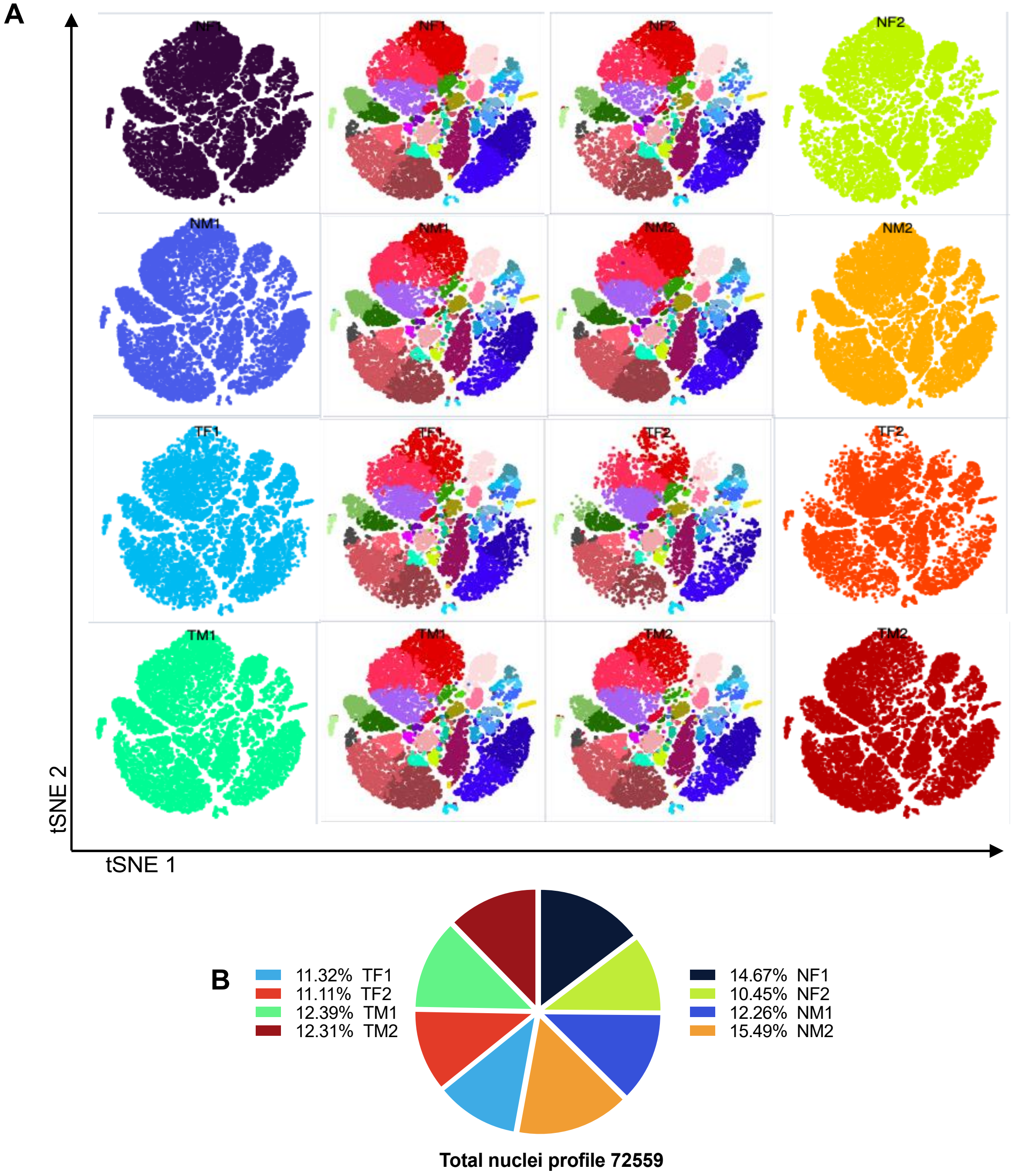


**Figure S6. Single-nucleus transcriptomic landscape of nTg and CCNA2-Tg mouse hearts. A)** t-SNE plot clustering the nuclei of nTg and CCNA2-Tg conditions (*n*=8 total). **B)** Pie chart representing the percentage of nuclei profile in all conditions of nTg and CCNA2-Tg mice. (*n*= 72,559 total nuclei)

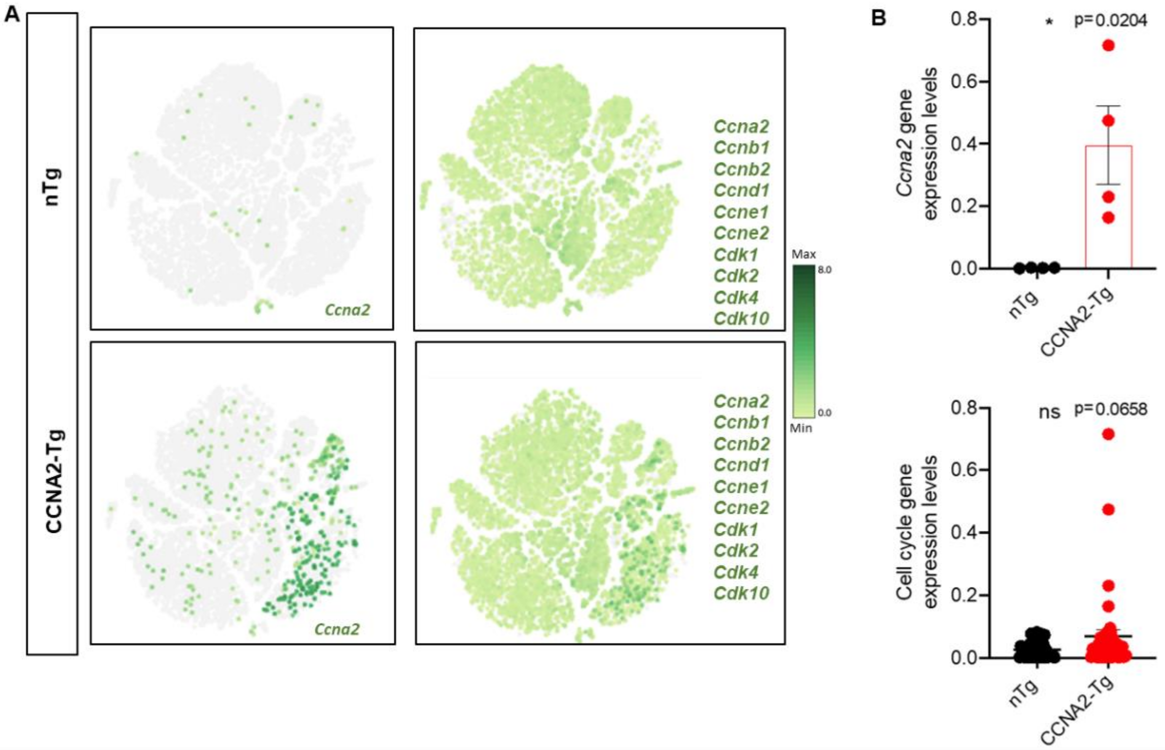


**Figure S7. CCNA2 and cell cycle gene expression across cardiac subclusters in nTg and CCNA2-Tg mice. A)** t-SNE plots and **B)** representative scatter plots of CCNA2 and cell cycle genes expression across the combined transcriptomic profiles of all subclusters of nTg and CCNA2-Tg mice. Bars represent mean ± s.e.m. Each point represents an individual value; the mean value is represented by the horizontal line. Error bars represent s.e.m.

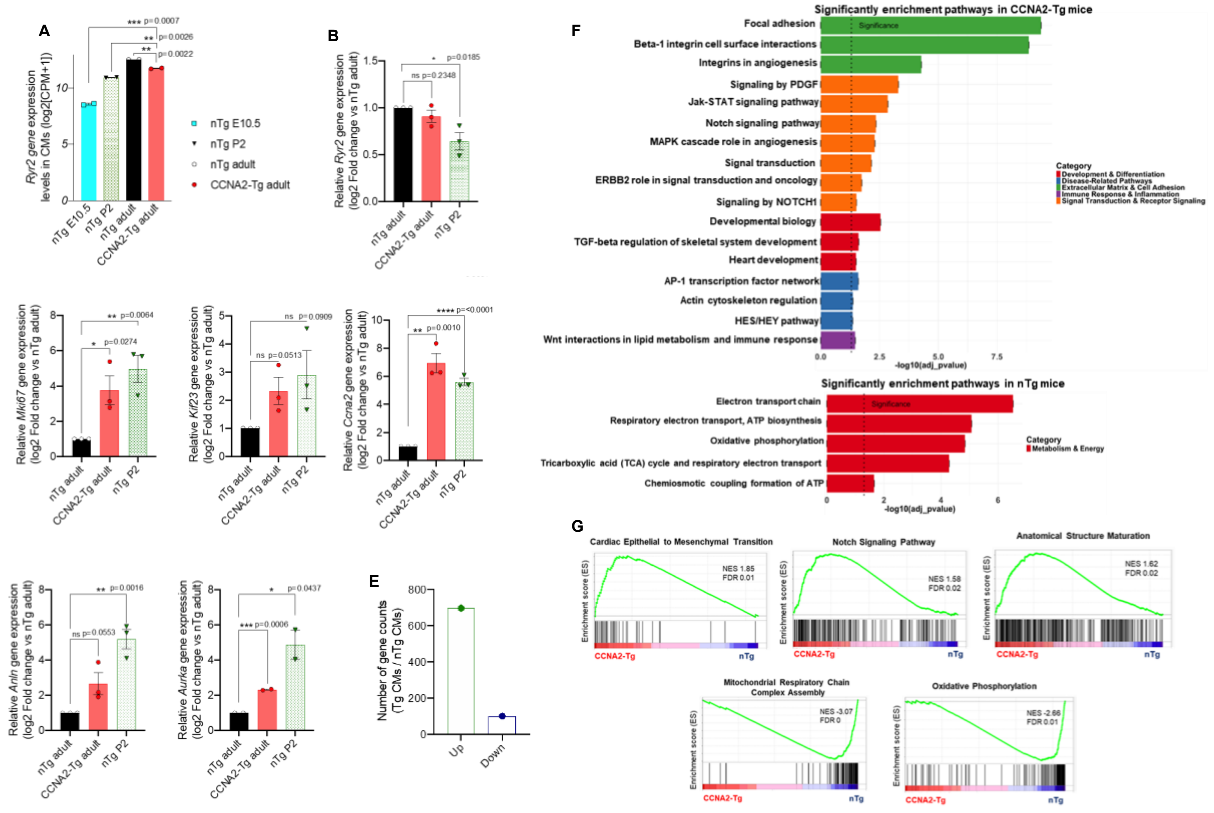

**Figure S8. Transcriptomic profiling reveals partial dedifferentiation and developmental gene program activation in CCNA2-Tg cardiomyocytes. A)** *Ryr2* gene expression levels across cardiomyocyte development stages, including embryonic (E10.5), postnatal (P2), adult nTg, and adult CCNA2-Tg mouse cardiomyocytes. **B)** RT-qPCR validation of *Ryr2* gene, **C)** proliferation, and **D)** cytokinesis-associated genes in P2, adult nTg and CCNA2-Tg cardiomyocytes. Expression values were normalized to *Gapdh* and are presented as fold change relative to adult nTg (mean ± s.e.m.); *n=3* independent experiments, with matched data from one experiment were excluded following ROUT outlier detection (Q=1%) applied uniformly across groups. **E)** Number of differentially expressed genes (Up or Down) in adult CCNA2-Tg mouse cardiomyocytes as compared to adult nTg mouse cardiomyocytes. F) Significantly enriched pathways in adult CCNA2-Tg and nTg mouse cardiomyocytes (P-value adjusted <0.05), analyzed using BioPlanet. **G)** Gene set enrichment analysis (GSEA) in CCNA2-transgenic versus nTg cardiomyocytesshowing a significant downregulation of oxidative phosphorylation and mitochondrial respiratory chain complex assembly pathways in CCNA2-transgenic cardiomyocytes, may suggest a metabolic shift consistent with dedifferentiation. Conversely, pathways such as cardiac epithelial-to-mesenchymal transition (EMT) and Notch signaling were upregulated, indicating activation of developmental and reprogramming pathways that facilitate a more regenerative state.

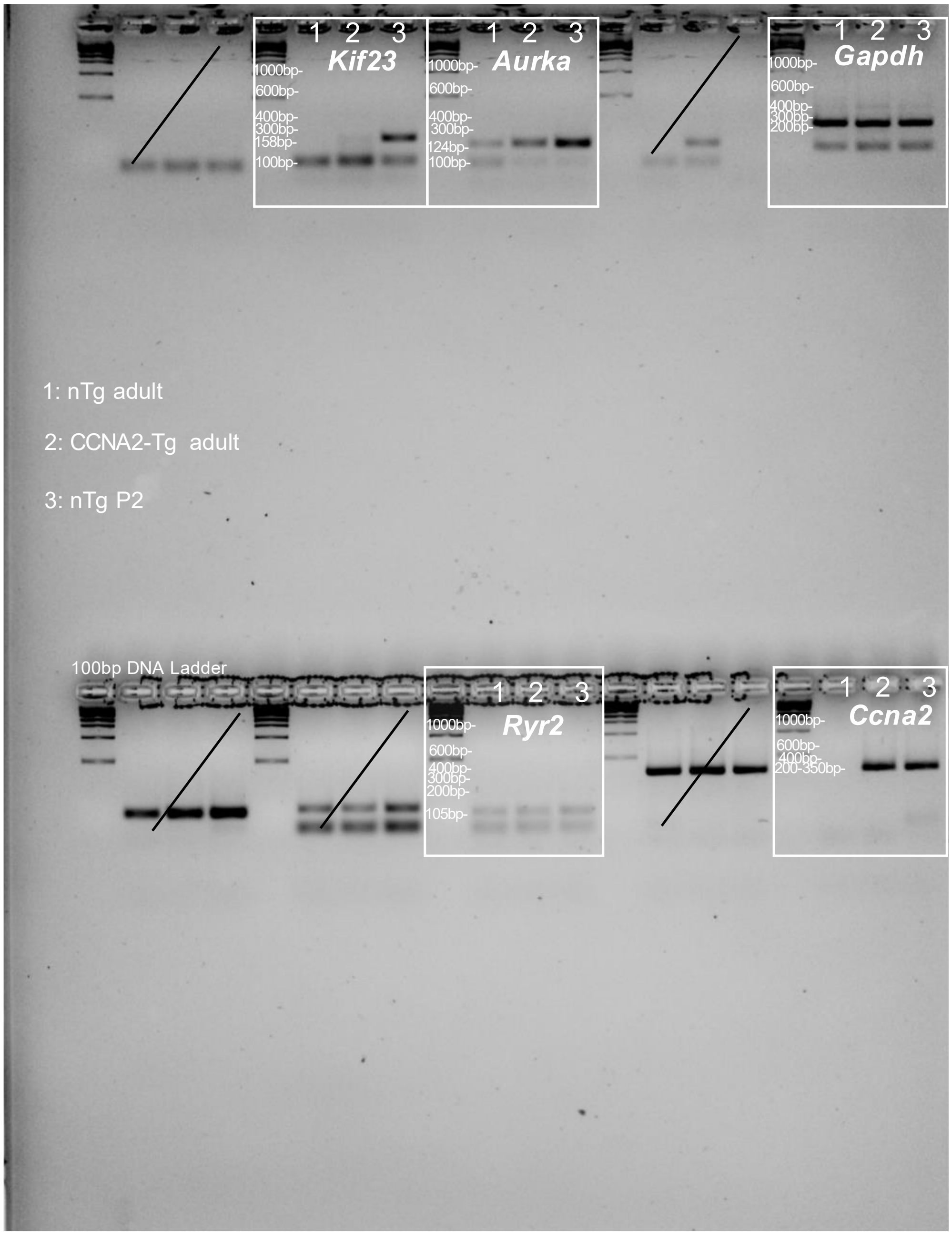


**Figure S9. Uncropped agarose gel electrophoresis of PCR products used in Figure S4F.** Full uncropped 2% agarose gel showing conventional PCR amplification of *Kif23, Aurka, Gapdh, Ryr2* and *Ccna2* in nTg adult (1), CCNA2-Tg adult (2) and P2 (3) samples used in Figure S4F. 100 bp DNA ladder (Thermo Fisher Scientific, Cat# 15628019) is included.

**Supplementary tables**
**Table S1: Primers sequences used in this study.**

| Gene | Forward primer sequence (5' - 3') | Reverse primer (5' - 3') |
|---|---|---|
| *Ryr2* | TTC CCC AAG ATG GTG ACA AG | CTC CAG CAG GTA GCT CAG GT |
| *Mki67* | GAG GAG AAA CGC CAA CCA AGA G | TTT GTC CTC GGT GGC GTT ATC C |
| *Kif23* | AGG CAT GGT GAA CAG AAA CGT CG | CTC TGG CTT CTC AGG TTT GGG T |
| *Ccna2* | CAC AAT CAC TTC TGA ATG TAG A | TAG AAG GAC ACC TAG TCA GAC AA |
| *Anln* | AGC ATC AGA CCT GGA GGT TGA G | CCT TCC TCT AGG ACG TCA CTG A |
| *Aurka* | TCA TCC TGG CTC TGA AGG TGC T | CCA TAC AGC CTG AGG ATG G |
| *Gapdh* | CCA GCT ACT CGC GGC TTT A | GTT CAC ACC GAC CTT CAC CA |

**Table S2: *The ARRIVE Essential 10 (Compliance Questionnaire).**

| Item | | Question(s) | Answers | |
|---|---|---|---|---|
| 1 | Study Design | Are all experimental and control groups clearly identified? | X | Yes, for at least one experiment<br>No |
| | | Is the experimental unit (e.g. an animal, litter or cage of animals) clearly identified? | X | Yes, for at least one experiment<br>No |
| 2 | Sample Size | Is the exact number of experimental units in each group at the start of the study provided (e.g. in the format 'n='? | X | Yes, for at least one experiment<br>No |
| | | Is the method by which the sample size was chosen explained? | X | Yes, for at least one experiment<br>No |
| 3 | Inclusion & Exclusion Criteria | Are the criteria used for including and excluding animal, experimental units, or data points provided? | X | Yes, for at least one experiment<br>No |
| | | Are any exclusions of animals, experimental units, or data points reported, or is there a statement indicating that there were no exclusions? | X | Yes, for at least one analysis<br>No |
| 4 | Randomization | Is the method by which experimental units were allocated to control and treatment groups described? | X | Yes, for at least one experiment<br>No |
| 5 | Blinding | Is it clear whether researchers were aware of, or blinded to, the group allocation at any stage of the experiment or data analysis? | X | Yes, for at least one experiment<br>No (X) |

| | | | | |
|---|---|---|---|---|
| 6 | Outcome Measures | For all experimental outcomes presented, are details provided of exactly what parameters were measured? | X | Yes, for at least one experiment<br><br>No |
| 7 | Statistical Methods | Is there statistical approach used to analyze each outcome detailed? | X | Yes, for at least one analysis<br><br>No |
| | | Is there a description of any methods used to assess whether data met statistical assumptions? | X | Yes, for at least one analysis<br><br>No<br><br>Not applicable |
| 8 | Experimental Animals | Are all species of animals used specified? | X | Yes, for at least one analysis<br><br>No |
| | | Is the sex of the animals specified to species? | X | Yes, for at least one experiment<br><br>No<br><br>Not applicable |
| | | Is at least one of the age, weight, or developmental stage of the animals specified? | X | Yes, for at least one experiment<br><br>No |
| 9 | Experimental Procedures | Are both the timing and frequency with which procedures took place specified? | X | Yes, for at least one experiment<br><br>No |
| | | Are details of acclimatization periods to experimental locations provided? | X | Yes, for at least one experiment<br><br>No |
| 10 | Results | Are descriptive statistics for each experimental group provided, with a measure of variability (e.g., mean and SD, or median and range)? | X | Yes, for at least one experiment<br><br>No<br><br>Not applicable to the type of data collected |
| | | Is the effect size and confidence interval provided? | X | Yes, for at least one experiment<br><br>No<br><br>Not applicable to the type of analysis used |

*Percie du Sert, N. et al. The ARRIVE guidelines 2.0: Updated guidelines for reporting animal research. PLoS Biol 18, e3000410 (2020).

**Supplementary Movies**

**Movie S1:** The real-time, live-imaging movie of human adult cardiomyocytes (55-year-old male) as described in **Figure 1**.
**Movie S2:** Real-time, live-imaging movie of human adult cardiomyocytes (41-year-old female) as described in **Figure 1**.

**Supplementary Data:** Supplementary Data contain the raw values and extended datasets supporting the Figures.
